# Mitochondrial cardiolipin metabolism controlled by tafazzin enables ferroptosis

**DOI:** 10.1101/2024.10.25.620299

**Authors:** Yvonne Wohlfarter, Judith Hagenbuchner, Utku Horzum, Gregor Oemer, Andreas Winter, Markus Seifert, Julian Schwärzler, Janik Kokot, Pablo Hernansanz Agustín, Viktorija Juric, Luiz Felipe Garcia Suoza, Heribert Talasz, José Antonio Enríquez Domínguez, Hesso Farhan, Timon E. Adolph, Günter Weiss, Johannes Zschocke, Markus A. Keller

## Abstract

Mitochondria are important producers of reactive oxygen species, which are involved in triggering ferroptosis, a lipid peroxidation driven form of cell death. Paradoxically, in the rare inherited metabolic disease Barth Syndrome, we discovered a protection from erastin-induced ferroptosis, despite intrinsically elevated mitochondrial ROS levels. The affected transacylase tafazzin, which is mutated in Barth Syndrome, is pivotal for remodeling of the dimeric phospholipid cardiolipin. They unique to mitochondria and essential for shaping their membrane functionalities. We investigated which downstream effects of the pathogenic membrane alterations are responsible for the protective effect against ferroptosis. We found that while iron metabolism, the unsaturation of membrane lipids, and the metabolic activity of the cells were modifying factors, they were not causal. However, we observed that cardiolipin abnormalities are not limited to impair only inner, but also outer mitochondrial membrane protein complexes. Specifically, they impact abundance and oligomerization of voltage-dependent anion channels (VDAC) in response to oxidative stress. We found that tafazzin deficiency via alteration of cardiolipins affects VDAC functionality, thereby modulating small molecule transport and signaling between mitochondria and the remaining cell. This is in line with a reduction of mitochondria-associated membranes (MAM) sites that are formed through VDACs and trapping ROS in mitochondria where they are unable to contribute to ferroptosis. These findings demonstrate that the mitochondrial membrane architecture impacting on subcellular small molecule distribution crucially impact on the manifestation of cell fate decisions, including ferroptosis.

## Introduction

Cell death mechanisms are fundamental for mammalian development, homeostasis, disease progression and prevention. Among the key players in these intricate processes, mitochondria have emerged as vital contributors. They exert their influence through their involvement in cellular reactive oxygen species (ROS) production, maintenance of redox balance, and possession of crucial metabolites and enzymes necessary for apoptosis, necroptosis, and pyroptosis (Nguyen et al., 2023). Moreover, they are increasingly recognized for their involvement in ferroptosis, an iron dependent type of regulated cell death characterized by lipid peroxide accumulation, which operates independently of caspase activation or DNA fragmentation (Yang and Stockwell, 2008; Dixon et al., 2012; Gao et al., 2019). Lipid peroxide accumulation in ferroptosis can be induced through well-established pathways, such as inhibition of glutathione peroxidase (GPx4) activity by substances like RSL3 or the depletion of glutathione (GSH), the cofactor of GPx4, via the inhibition of the cysteine-glutamate transporter system (system x_c_) by erastin (Yang and Stockwell, 2008; Dixon et al., 2012).

Despite being a prototypical ferroptosis inducer, erastin exerts multifaceted cellular effects by additionally targeting voltage-dependent anion channels (VDAC) in the outer mitochondrial membrane (OMM), reversing the tubulin blockage and thereby changing VDAC conductance (Yagoda et al., 2007; Yang and Stockwell, 2008; Dixon et al., 2012). VDACs are crucial for the exchange of a broad range of ions and small molecules (Bayrhuber et al., 2008) and serve as major escape routes of ROS (Han et al., 2003). Besides that, VDACs are essential for the formation of mitochondria-associated membranes (MAMs), which are established through interactions between ER-localized inositol-1,4,5-triphosphate receptors (IP_3_R) with VDACs (Szabadkai et al., 2006; Peruzzo et al., 2020; Zhang et al., 2024).

The structural integrity and functional dynamics of mitochondrial transport systems are tied to the complex architecture of the embedding membranes. To meet their unique functional demands, mitochondrial membranes contain a dimeric phospholipid (PL) known as cardiolipin (CL). CLs are characterized by four tissue-specific, mostly highly unsaturated fatty acyl side chains (Oemer *et al*., 2018), which is generated by a remodeling process that is largely catalyzed by the transacylase tafazzin (Róg et al., 2009; Horvath and Daum, 2013; Basu Ball, Neff and Gohil, 2018). These lipids play a critical role in alleviating curvature stress induced by protein crowding in oxidative phosphorylation (OXPHOS) systems, due to their intrinsic negative curvature (Xu *et al*., 2019). CLs are vulnerable to lipid peroxidation, as they are a known binding partner of electron transport chain (ETC) complexes (Pfeiffer *et al*., 2003), the primary sources of ROS in cells (Hernansanz-Agustín and Enríquez, 2021), and harbor highly unsaturated side chains. Increased lipid peroxidation is vital for ferroptosis induction (Dixon *et al*., 2012). Mutations in the *TAFAZZIN* gene (OMIM #300394) lead to Barth Syndrome (OMIM #302060), an X-linked genetic disorder characterized by cardiomyopathy, skeletal muscle myopathy, growth delay and neutropenia (Barth *et al*., 1983; Bolhuis *et al*., 1991; Bione *et al*., 1996; Vreken *et al*., 2000).

In this study, we investigated the functional interplay between mitochondrial membrane lipid composition with a focus on CLs, and their role in the context of ferroptosis. We made the unexpected observation that dysfunctional CL remodeling led to resistance against erastin-induced ferroptosis, despite strongly elevated mitochondrial ROS levels. This counterintuitive effect could be traced back to a newly discovered mechanism by which CL impact the oligomerization of mitochondrial membrane-localized small molecule transporter systems. This novel insight into the interplay between the mitochondrial membrane architecture, mitochondrial signaling, and ferroptosis execution expands our understanding of how lipid-metabolic enzymes can intricately modulate cell death pathways across cellular compartments.

## Results

### Tafazzin-deficiency protects cells from ferroptosis

The primary consequences of a tafazzin-deficiency are pronounced cardiolipin (CL) abnormalities, characterized by CL depletion, monolyso-CL (MLCL) accumulation, and a more saturated CL profile (Figure 1A). Tafazzin-deficiency exclusively alters CL side chains and did not substantially impact other phospholipids (PL) (Oemer *et al*., 2021), which we additionally confirmed here in a subcellular manner (Figure 1B and Supplementary Figures 1 and 2). In tafazzin-deficient HEK 293T cells (ΔTAZ) CL abnormalities result in ∼4 fold increased lipid peroxidation levels (Figure 1C) and in 4-6 fold increased oxidative stress levels in mitochondria compared to controls (Ctrl, Figure 1D). Notably, most of the observed Bodipy 581/591 C11 (both oxidized and non-oxidized) accumulates in the ER (Supplementary Figure 3), indicating cell-wide phenotypic consequences derived from localized changes in inner mitochondrial membrane lipids. Additionally to increased oxidative stress levels, ΔTAZ cells show a strongly fragmented mitochondrial morphology (Figure 1E), which is in line with previous observations (He *et al*., 2013; Zhu *et al*., 2021; Komaragiri *et al*., 2022).

**Figure 1:**
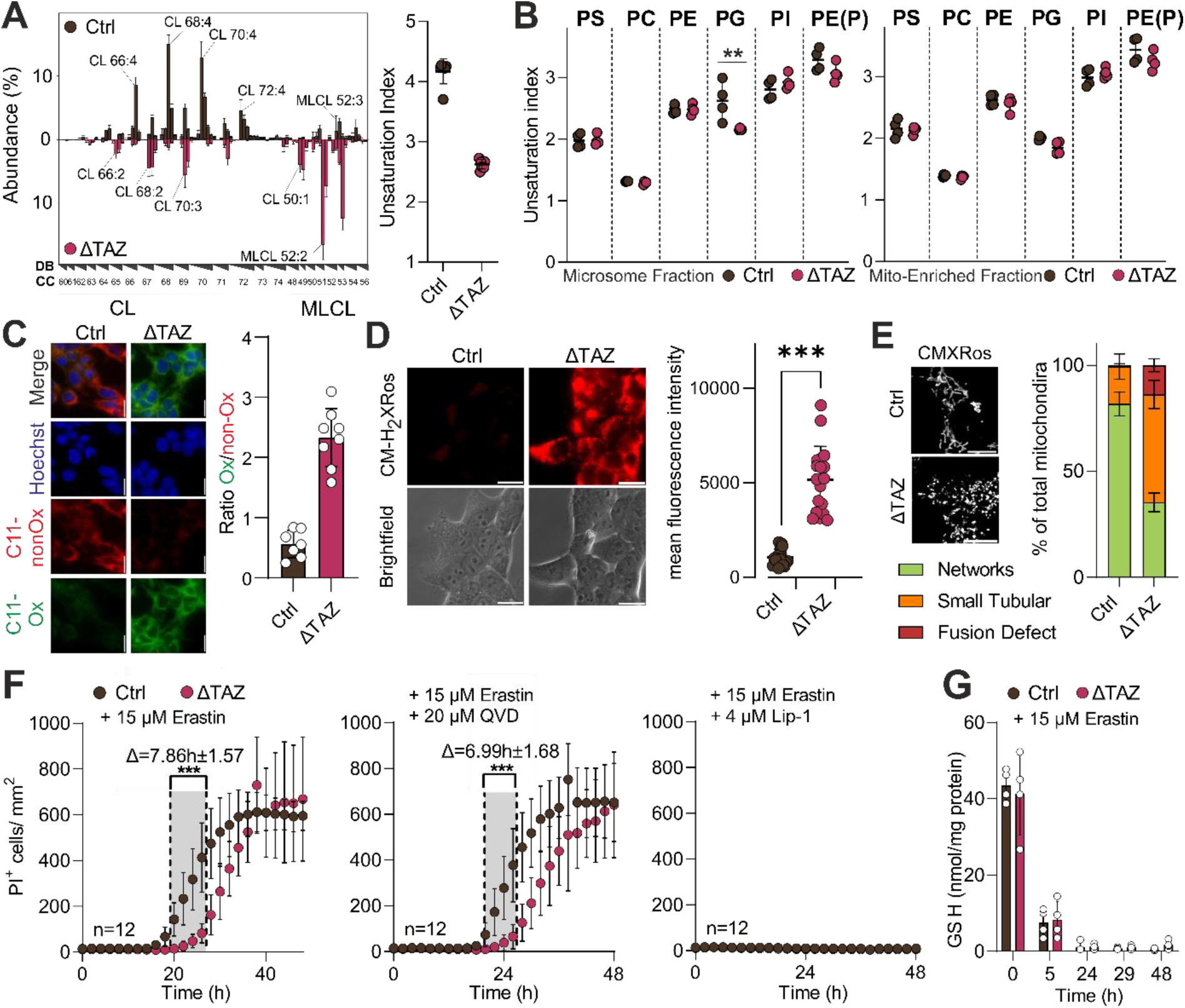
Tafazzin-deficiency protects against ferroptosis. A) Left: Relative abundance of CL species in Ctrl and ΔTAZ HEK 293T cells. Profiles are shown as mean ± SD and are normalized to total CL. CL species are indicated by their total number of side chain carbon atoms (CC) and organized according to the number of double bonds (DB). Right: Unsaturation index of CLs in Ctrl and ΔTAZ was calculated as weighted average and data is shown as mean ± SD. B) Unsaturation index of phospholipid species (phosphatidylserines (PS), phosphatidylcholines (PC), phosphatidylethanolamines (PE), phosphatidylglycerols (PG), phosphatidylinositols (PI), PE-plasmalogens (PE(P)) in Ctrl and ΔTAZ was calculated as weighted average and data is shown as mean ± SD. C) Live cell imaging of lipid peroxidation levels in Ctrl and ΔTAZ cells using Bodipy581/591 C11 (Bar size 20 µm). D) Mitochondrial ROS levels of Ctrl and ΔTAZ were analyzed by CM-H2XRos staining (reduced, 300 nM, Bar size 10 µm), (Unpaired t-test, p<0.001). E) Mitochondrial morphology stained with CMXRos (oxidized Mitotracker, 500 nM). Shown is the mean of three independent experiments; for each experiment 30–40 cells were analyzed. F) Cell death induction in Ctrl and ΔTAZ with 15 µM erastin followed by live cell imaging over 48h or 4 µM RSL3 over 4h, cell death was indicated via PI^+^ (1 µg/ml), (Ordinary one-way ANOVA, multiple testing corrected, p<0.001). G) GS-NEM depletion was measured over 48 hours after the start of a 15 µM erastin treatment in Ctrl and ΔTAZ cells using LC-MS/MS.

Surprisingly, even though these observed oxidative characteristics are expected to accelerate ferroptosis (Dixon *et al*., 2012; Doll *et al*., 2017), we observed an unexpected protective effect in cells with a tafazzin-deficiency. When treated with erastin, ΔTAZ cells exhibited a substantial and significant protection against ferroptotic cell death (Δ7.86 h ±1.57 h) (Figure 1F), which was further confirmed utilizing the ferroptosis and apoptosis inhibitors liproxstatin-1 (Lip-1) and Q-VD-Oph (QVD), respectively. Erastin-induced ferroptosis followed the previously reported wave like dying behavior (Riegman *et al*., 2020) in Ctrl, as well as ΔTAZ cells (Supplementary Figure 4), and we observed the expected dependence on cell density (Supplementary Figure 5) (Yang *et al*., 2019).

We did not observe differences in glutathione (GSH) metabolism that could explain this behavior. However, despite changes in the baseline GPx4 expression in ΔTAZ cells (Supplementary Figure 6A), steady-state level and erastin-induced depletion kinetics of GSH were unaffected (Figure 1G and Supplementary Figure 6B). Similarly, the tafazzin-dependent protective phenotype cannot be diminished by the addition of tetrahydrobiopterin (BH_4_), a known ferroptosis-counteracting antioxidant (Supplementary Figure 7) (Kraft *et al*., 2020). Furthermore, in Ctrl cells erastin induced mitochondrial fragmentation, which was unchanged in ΔTAZ cells that already showed fragmentation before the addition of erastin (Supplementary Figure 8).

As a first step to explain the observed protection conferred by tafazzin-deficiency, we modulated cellular iron availability and monitored its effect on the ferroptotic behavior. The addition of iron (+Fe) accelerated erastin-induced cell death (Figure 2A). However, despite the earlier onset of cell death in iron-saturated conditions, the protective effect of tafazzin-deficiency persisted. Moreover, tafazzin-deficiency offers such robust protection against ferroptosis that iron chelation with deferiprone (+DFP) does not further enhance this protective effect. To examine, whether ΔTAZ cells exhibit an alteration in iron metabolism, we quantified transporter systems involved in iron homeostasis (Figure 2B & Supplementary Figure 9). Despite some differences in TfR1 and IRP2, Ctrl and ΔTAZ cells showed no differences in the availability of their ferroptosis-critical labile iron pools (LIP) (Figure 2C&D), as well as in total iron levels (Figure 2E). This is also reflected in the unchanged activity of the mitochondrial iron-sulfur cluster-dependent enzyme aconitase (Figure 2F).

**Figure 2:**
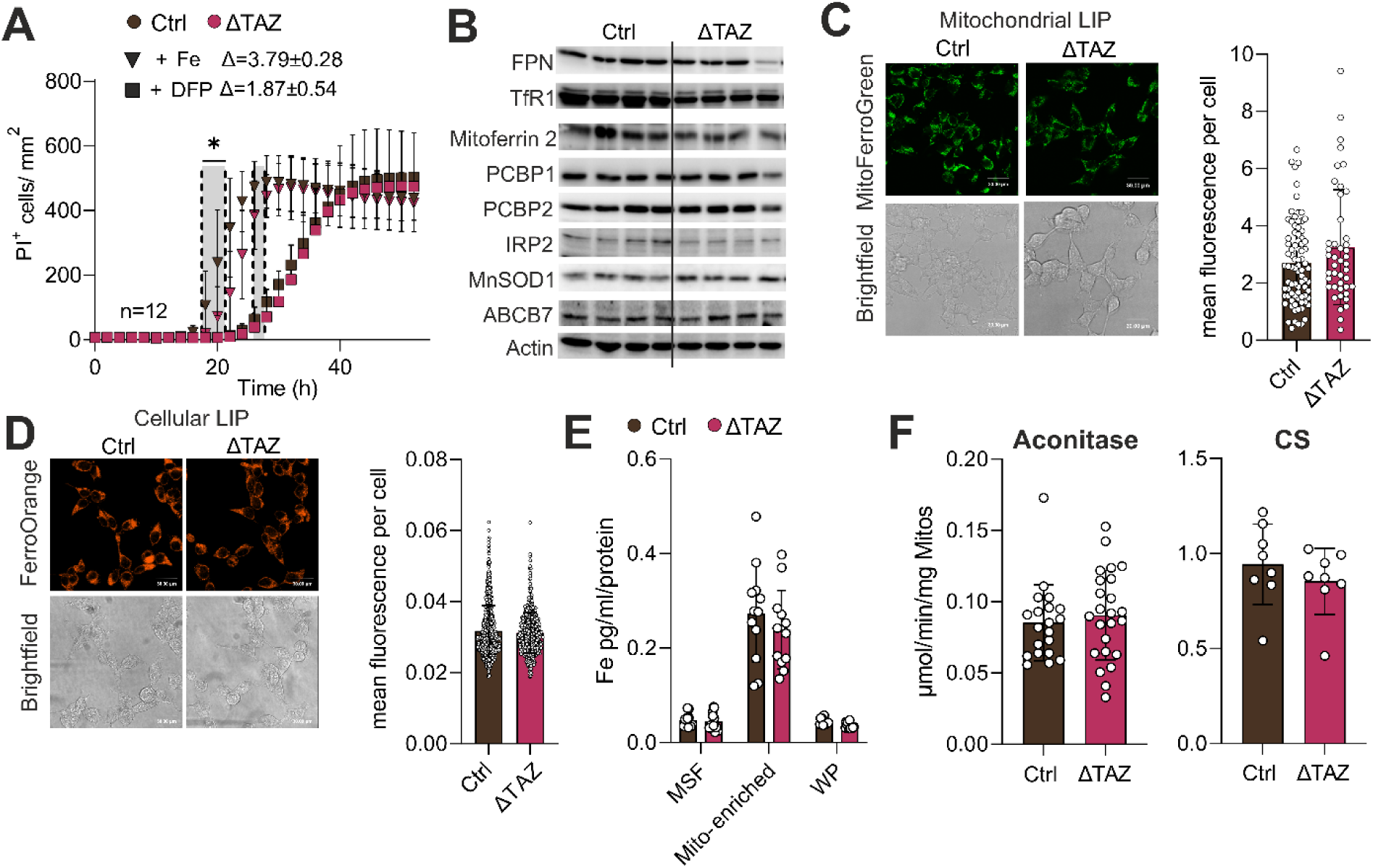
Iron availability is modulating ferroptosis, but does not explain the protection of a tafazzin-deficiency. A) Ferroptotic cell death kinetics in iron-supplemented (50 µM FeCl_2_), or iron-chelated (deferiprone (DFP)) conditions in Ctrl and ΔTAZ cells following exposure to 15 µM erastin. Live cell imaging was performed, and cell death was assessed using PI^+^ (1 µg/ml). Statistical analysis revealed significant differences between treatment conditions (Two-way ANOVA, multiple testing corrected, p=0.012). B) Ferroportin (FPN), Transferrin receptor 1 (TfR1), Iron regulatory protein 2 (IRP2), mitochondrial superoxide dismutase (MnSOD), and Poly C Binding Protein 1/2 (PCBP1/PCBP2), mitochondrial sodium dismutase (MnSOD1) and ATP binding cassette subfamily B member 7 (ABCB7) were measured via western blotting in Ctrl and ΔTAZ cells, across four different tafazzin knockouts and four different controls. Among these proteins, only the transferrin receptor (TfR1) was significantly lower expressed in ΔTAZ cells), (Two-way ANOVA, multiple testing corrected, p<0.001). C&D) Fluorescent quantification of mitochondrial and cytosolic labile iron pool (LIP) using FerroOrange and MitoFerroGreen according to manufacturer instructions. Intensity was analyzed using CellProfiler 4.2.6 (Stirling et al., 2021). E) Total iron quantification using atom absorption spectroscopy in the microsomal fraction (MSF), Mito-enriched and unfragmented cell fraction. Experiment was performed 3 individual times using 4 biological replicates each (n=12). F) Mitochondrial aconitase activity quantification and citrate synthase (CS) quantification in isolated mitochondria.

### The ferroptotic behavior cannot be explained by differences in CL saturation levels

To characterize the contribution of the lipid environment and CL unsaturation to the susceptibility of cells towards ferroptosis, we supplemented the cells with the poly-unsaturated linoleic acid (LA, FA18:2) and the saturated palmitic acid (PA, FA16:0), respectively. Especially the LA treatment strongly altered CL patterns and the related unsaturation index (Figure 3A) at constant growth rates (Supplementary Figure 10). However, supplemented fatty acids change the unsaturation and oxidizability of the entire lipidome (Oemer *et al*., 2021). Consequently, we observed that PA rich conditions delayed, while LA accelerated erastin-induced ferroptosis (Figure 3B, controls in Supplementary Figure 11). However, in both environments a clear protective effect of tafazzin-deficiency persisted.

**Figure 3:**
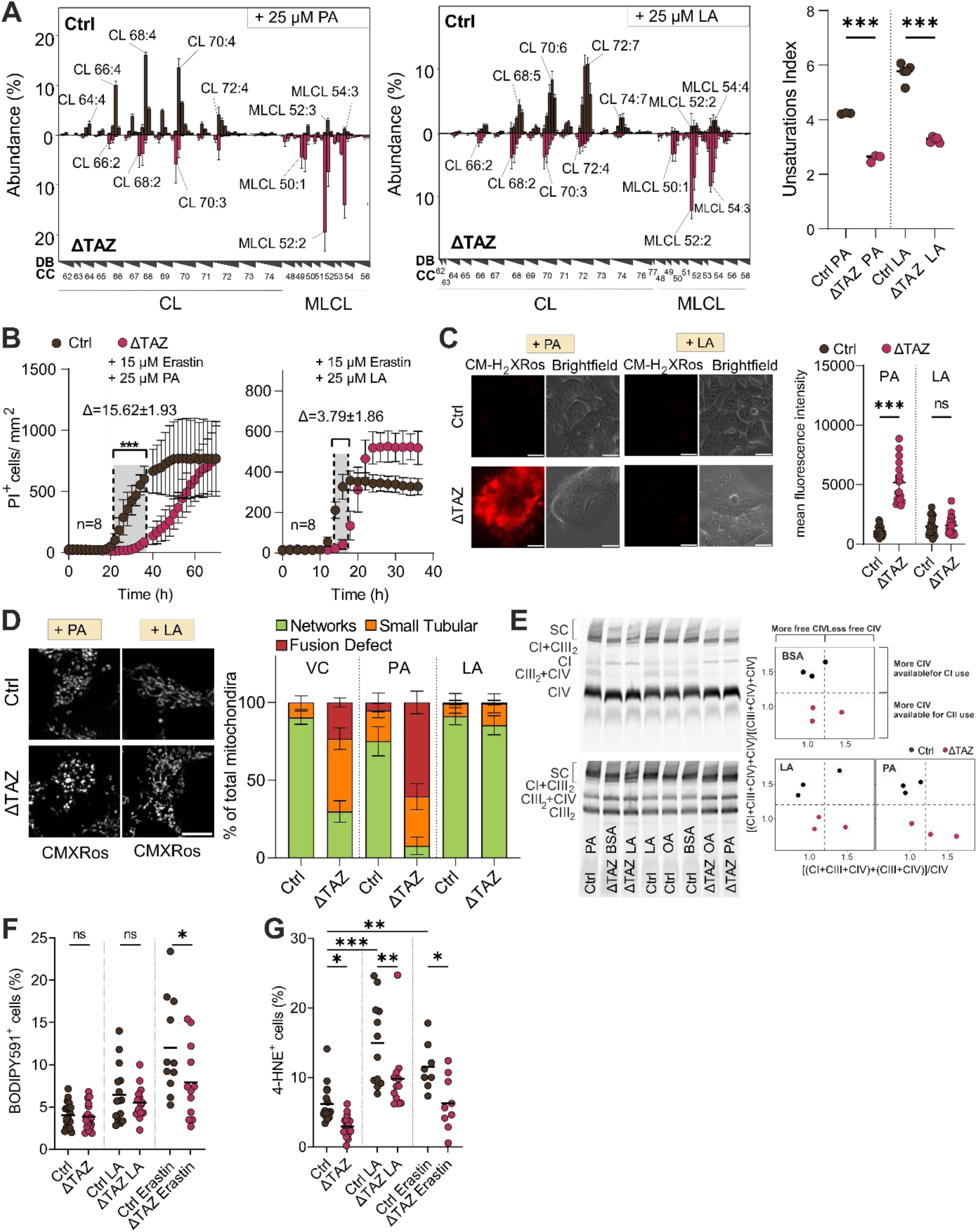
Lipid supplementation influences ferroptosis, mitochondrial morphology and function. A) Relative abundance of CL species in Ctrl and ΔTAZ supplemented with 25 µM linoleic (LA) or palmitic acid (PA). Profiles are shown as mean ± SD and are normalized to total CL. CL species are indicated by their total number of side chain carbon atoms (CC) and organized according to the number of double bonds (DB). B) Cell death induction in Ctrl and ΔTAZ with 15 µM erastin in combination with 25 µM linoleic acid (LA) or 25µM palmitic acid (PA) followed by live cell imaging over 72h, cell death was indicated via PI^+^ (1 µg/ml), (Two-way ANOVA, multiple testing corrected, p<0.001). C) Mitochondrial ROS levels of Ctrl and ΔTAZ in combination with 25 µM LA or 25 µM PA were analyzed by CM-H_2_XRos staining (300 nM, Bar size 10 µm), (Ordinary one-way ANOVA, multiple testing corrected, p<0.001). D) Mitochondrial morphology of Ctrl and ΔTAZ in combination with 25 µM linoleic acid (LA) or 25 µM palmitic acid (PA) stained with CMXRos (500 nM). Shown is the mean of three independent experiments; for each experiment 30–40 cells were analyzed. E) Supercomplex analysis in Ctrl and ΔTAZ supplemented with 25 µM LA or 25 µM PA. F) FACS analysis of lipid peroxidation in Ctrl and ΔTAZ supplemented with 25 µM of LA or after 8h of erastin treatment. ΔTAZ showed significantly lower levels of lipid peroxidation compared to control cells (Ordinary one-way ANOVA, multiple testing corrected, p= 0.0118). G) FACS analysis of 4-hydroxynonenal (4-HNE) levels in Ctrl and ΔTAZ supplemented with 25 µM of LA or after 8h of erastin treatment. ΔTAZ showed significantly lower levels throughout tested conditions (Ordinary one-way ANOVA, multiple testing corrected, *p<0.033, **p<0.002, ***p<0.001).

Interestingly both, ROS levels and mitochondrial morphology in ΔTAZ cells, were reduced to levels observed in Ctrl cells upon LA supplementation (Figure 3C&D). In contrast, the PA treatment did not alter ROS formation and even deteriorated mitochondrial morphology. Despite restored morphology and ROS levels in LA-treated ΔTAZ, the protective effect of tafazzin-deficiency against ferroptosis remained (Figure 3B).

Malfunctions in the supercomplex assembly of OXPHOS complexes is one of the main molecular features of Barth Syndrome (McKenzie *et al*., 2006). Thus, we next investigated its association with the observed behavior of the mitochondrial lipid environment, ROS levels, and morphology. Indeed, ΔTAZ cells were impaired in forming higher molecular mitochondrial supercomplexes (Figure 3E). Supercomplexes can exist in various configurations such as CI/CIII/CIV, CI/CII or CIII/CIV (Mileykovskaya and Dowhan, 2014). The ratio between different configurations shows that specifically Complex IV availability shifts from CI towards CII in ΔTAZ cells. However, this phenotype was not altered by any of the tested lipid environments (see Figure 3E right panels). Under physiological conditions, this appears to be compensated, as we did not record any significant differences in the respiratory capacity in NADH-dependent (N), succinate-dependent (S) and N/S coupling control states (Supplementary Figure 12).

To assess further consequences of linoleic acid and erastin treatment, we next measured lipid peroxidation levels and a resulting byproduct, 4-hydroxynonenal (4-HNE). ΔTAZ cells exhibited significantly lower levels of oxidized Bodipy C11 upon erastin treatment compared to controls (Figure 3F) and, remarkably, despite elevated baseline mitochondrial ROS, ΔTAZ cells displayed reduced 4-HNE levels under all tested conditions (Figure 3G). This suggests that erastin-induced ferroptosis, in which mitochondria play a pivotal role (Gao *et al*., 2019), appears to be less effective in signaling its effects in the context of a tafazzin-deficiency.

### Impairment of ER-mitochondrial signaling protects against ferroptosis in a VDAC- dependent manner

A mitochondria-centered regulatory mechanism in tafazzin-deficient cells is further substantiated by the finding that no ferroptosis protection was observed when cell death was triggered with the GPx4 inhibitor RSL3 (Figure 4A). Erastin is known to interact with the VDACs in the OMM and changing their conductibility (Figure 4B). To examine if the protection is caused by VDAC dysregulation, we monitored VDAC1/3 expression levels during ferroptosis at 0, 8, 16, and 24 hours post-erastin treatment. In control cells, a considerable increase in VDAC1/3 expression is induced within the first 8 hours (Figure 4C), suggesting a regulatory involvement of VDACs in the early ferroptotic cascade. Interestingly, already in the untreated state a trend towards lower VDAC1/3 expression in ΔTAZ cells was observable. This effect became more pronounced and significant after 8 hours, showing that ΔTAZ cells lack the capacity to efficiently upregulate VDAC1/3. In contrast, expression of the mitochondrial calcium uniporter (MCU) was unaltered throughout ferroptosis (Figure 4C, Supplementary Figure 13). VDAC abundance and their oligomerization state have been shown to be crucial factors for ferroptosis susceptibility (Zhang *et al*., 2024). Therefore, we employed two VDAC oligomerization inhibitors, Vbit4 and NSC, to investigate their individual impacts on erastin- induced ferroptosis (Figure 4D). Our results demonstrate an extensive protection against ferroptosis in both Ctrl and ΔTAZ cells as a consequence of the oligomerization impairment, which was further confirmed utilizing the ferroptosis and apoptosis inhibitors liproxstatin-1 (Lip- 1) and Q-VD-Oph (QVD), respectively (Supplementary Figure 14).

**Figure 4:**
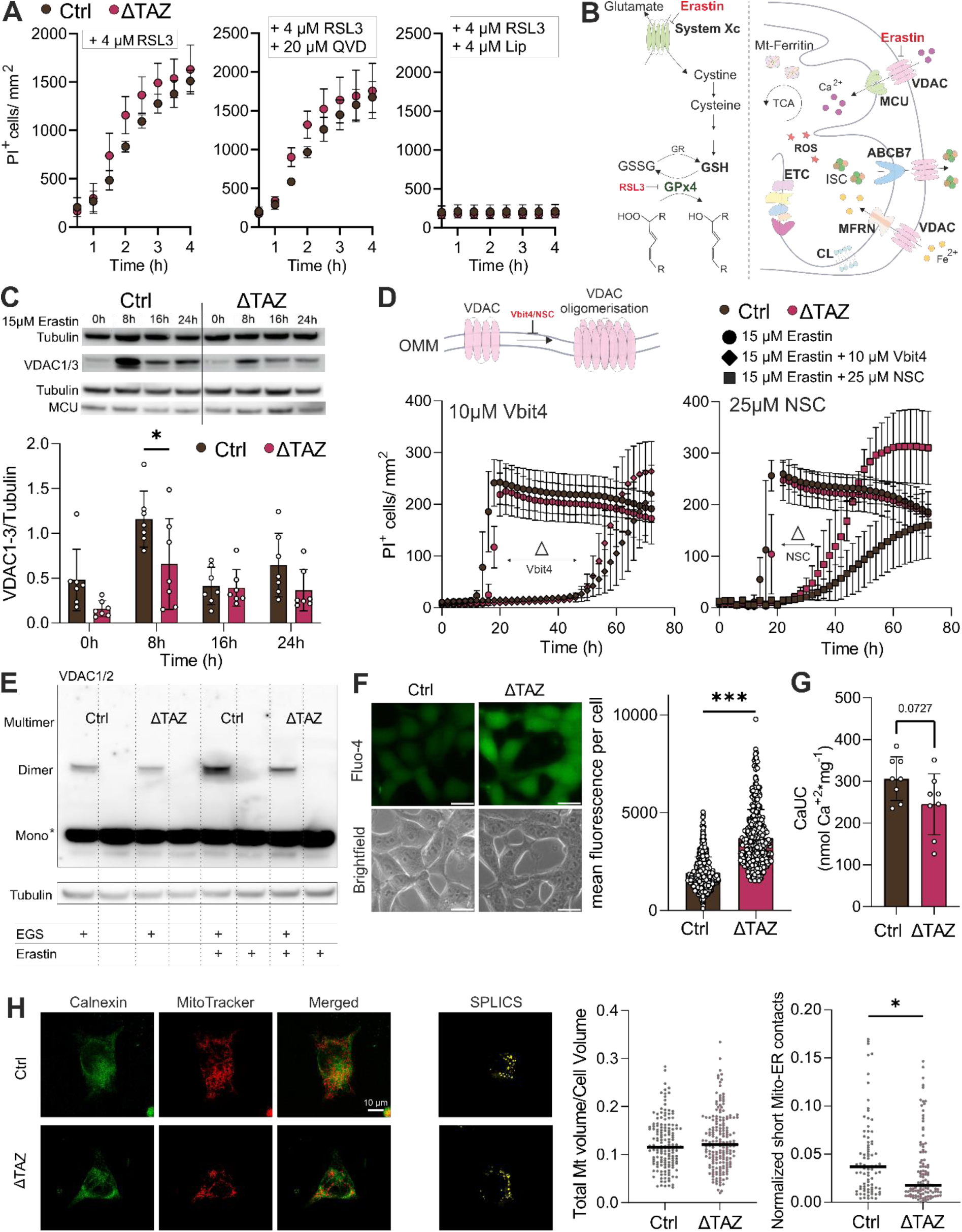
Abnormal CLs reduce mitochondria-ER contact sites and alter VDAC dynamics during ferroptosis. A) Cell death induction in Ctrl and ΔTAZ with 4 µM RSL3 followed by live cell imaging over 48h or 4 µM RSL3 ± 20 µM QVD ± 4 µM Lip-1 over 4h, cell death was indicated via PI^+^ (1 µg/ml). B) Schematic representation of cellular pathways that influence ferroptosis. Abbreviations: ATP binding cassette subfamily B member 7 (ABCB7), Calcium (Ca^2+^), Cardiolipin (CL), electron transport chain (ETC), glutathione peroxidase 4 (GPx4), Glutathione (GSH), oxidized GSH (GSSG), Iron (Fe), Iron sulfur cluster (ISC), Mitochondrial calcium uniporter (MCU), Mitoferrin (MFRN), Reactive oxygen species (ROS), Tricarboxylic acid cycle (TCA), Voltage dependent anion channel (VDAC). C) VDAC1/3 and MCU levels were assessed before and after treatment with 15 µM erastin using western blotting. Measurements were conducted at 8h, 16h, and 24h following treatment initiation (Two-way ANOVA, multiple testing corrected, p=0.014). D) Cell death induction in Ctrl and ΔTAZ with 15 µM erastin combined with 10 µM Vbit4 and 25 µM NSC respectively followed by live cell imaging over 72h, cell death was indicated via PI^+^ (1 µg/ml). E) Evaluation of EGS cross-linked VDAC multimers before and after 8h of 15 µM erastin treatment. Ctrl cells exhibited higher multimer levels in untreated conditions compared to ΔTAZ, which were further upregulated upon erastin treatment, whereas ΔTAZ cells failed to show such upregulation. F) Baseline calcium levels of Ctrl and ΔTAZ were analyzed by Fluo-4 staining according to manufacturer’s conditions (Bar size 20 µm) (Unpaired-t-test p<0.001). G) Calcium uptake capacity (CaUC) was measured using the O2k (Oroboros Instruments, Innsbruck) ΔTAZ could take up less calcium before mitochondrial burst than controls (Unpaired-t-test p=0.0727). H) ER-Mitochondria interactions using a SPLICS Mt-ER-Short P2A plasmid, ΔTAZ cells show fewer contact sites, but encompass the same total mitochondrial volume normalized to cell volume (Unpaired t-test p=0.13).

To further validate this effect, we chemically cross-linked VDACs with the cross-linking agent EGS (ethylene glycol bis(succinimidyl succinate)) and revealed that ΔTAZ cells have reduced levels of dimeric and multimeric VDAC1/2 oligomers compared to controls (Figure 4E, additional blots in Supplementary Figure 15), mimicking the CL-dependent supercomplex assembly defect of the respiratory chain shown in Figure 3E. Thus, ΔTAZ cells are unable to sufficiently upregulate VDAC expression and also fail to induce VDAC oligomerization during the early stages of erastin-dependent ferroptosis.

Metabolically, VDACs are well-characterized for mediating calcium transport between the cytosol and mitochondria (Figure 4B). To investigate whether the observed changes in VDAC behavior are significant enough to have respective functional consequences, we proceeded to measure cytosolic calcium levels (Figure 4F). Indeed, we observed a cytosolic calcium accumulation in combination with a reduced calcium uptake capacity of mitochondria in ΔTAZ cells (Figure 4F&G), indicating general disruptions of metabolic communication between mitochondria and the remaining cell as a consequence of tafazzin-deficiency.

Furthermore, VDACs play a central role in establishing mitochondria-associated membranes (MAMs), which are a hub for molecular exchange between ER and mitochondria. Recently, it has been demonstrated that deficient MAMs can impact on the susceptibility to ferroptosis (Zhang *et al*., 2024). Therefore, we investigated whether the VDAC abnormalities observed in the BTHS-model are affecting MAMs and thus contribute to the observed protection against ferroptosis. We characterized ER-mitochondria interactions using a split-GFP system. Indeed, we observed significantly reduced contact site numbers in the ΔTAZ model compared to controls with the relative mitochondrial volume remaining constant (Figure 4H).

## Discussion

The role of mitochondria in ferroptosis is described as complex and highly context dependent (Gao *et al*., 2019). As one of the primary sources of ROS, mitochondrial metabolism is often associated with the rapid depletion of GSH, which is required as a GPx4 cofactor (Dixon *et al*., 2012; Yang *et al*., 2014), and the accumulation of lipid-ROS in a pro-ferroptotic manner (Dixon *et al*., 2012). Paradoxically, in our Barth Syndrome cell model, we observed a protection against erastin-induced ferroptosis (Figure 1F), despite having inherently high levels of mitochondrial ROS (Figure 1D) and increased fission (Figure 1E). We showed that this finding can be explained by a cardiolipin aberration-induced impairment of VDAC oligomerization and expression (Figure 4). This could lead to the effect that instead of being readily released to the cell, a portion of ROS remains trapped in mitochondria, where they accumulate and cannot further contribute to the execution of ferroptosis.

In a similar manner - and likely caused by the same underlying functional effects - we find that the transport of calcium ions is impaired in tafazzin-deficiency, in our model (Figure 4F&G) and others (Ghosh *et al*., 2020, 2021). This demonstrates that the exact membrane cardiolipin composition can significantly contribute to the regulation of cell fate decisions. Such an effect is clearly not limited to Barth Syndrome, but should also be considered in other pathological conditions that impact on mitochondrial membrane constitutions.

The cardiolipin abnormalities in Barth Syndrome intrinsically point towards a mitochondrial origin of the protective effect against ferroptosis. This is further substantiated by the fact that the phenotype is present only in erastin-, but not RSL3-induced (Figure 4A) ferroptosis. Ferroptosis has been reported to coincide with VDAC depletion (Yang *et al*., 2020) and with the induction of VDAC oligomerization (Zhang *et al*., 2024). Both phenomena are observed in our model system (Figure 4). However, we find that VDAC expression is first upregulated during early ferroptosis in controls, a process that tafazzin-deficient cells are unable to efficiently conduct (Figure 4C). At time points closer to ferroptosis onset a degradation of VDACs is observed. A reduction of VDAC expression has been linked to a decreased ferroptosis susceptibility (Zhang *et al*., 2024), aligning to the here described protective effect against ferroptosis in tafazzin-deficiency (Figure 1F). Furthermore, the assembly of VDAC- multimers appears to be an important step in ferroptosis execution, as erastin triggered the induction of VDAC oligomerization in control cells (Figure 4E). In line with this, we demonstrated that blocking VDAC oligomerization with specific inhibitors rendered cells highly resistant to ferroptosis (Figure 4D). In contrast to controls, tafazzin-deficient cells were unable to efficiently induce VDAC oligomerization, which strongly resembles the well-described disruption of respirasomes (Figure 3E) (McKenzie *et al*., 2006; Chatzispyrou *et al*., 2018).

VDACs have been reported as influential in other forms of regulated cell death. For instance, increased VDAC oligomerization and abundance have been associated with apoptotic susceptibility (Abu-Hamad *et al*., 2009). Tafazzin-deficient cells have been shown to fail in activating the intrinsic apoptosis pathway, due to the lack of CLs in the OMM (Sorice *et al*., 2004). Although this resistance seems to be primarily caused by the missing CL - caspase-8 binding platform (Gonzalvez *et al*., 2008), preventing BID cleavage and consequently tBid/Bax oligomerization (Kuwana *et al*., 2002).

A core question is how CL, which is mainly enriched in the IMM, can influence the properties of the OMM in a way that it affects VDAC dynamics (Ardail *et al*., 1990). VDACs are primarily localized at contact sites between the IMM and OMM, where the OMM CL content increases from 4% to 20% (Ardail *et al*., 1990), reaching levels comparable to those in the IMM. Independent of this, it also has been shown that already relatively moderate CL concentrations in the OMM can impact on the oligomerization behavior of protein complexes (Kuwana *et al*., 2002) and modulate membrane packing stress (Rostovtseva *et al*., 2006). Of note, tafazzin-deficiency not only causes CL depletion but also a substantial accumulation of MLCLs (Figure 1A). The individual contributions of these lipid species to the observed VDAC behavior cannot be disentangled. For instance, while it has been shown that CL abundance is inversely correlated with VDAC oligomerization, the potential additional effects of MLCL have not been considered (Betaneli, Petrov and Schwille, 2012). This uncertainty aligns with an ongoing scientific debate about which CL characteristics are primarily causative for the Barth Syndrome phenotype (Malhotra *et al*., 2009).

VDACs play a dual role in MAMs: through the interaction with IP_3_R, which stabilizes the MAM environment, and by functioning as a small molecule transport system. VDACs are implicated in the exchange of small molecules and ions such as lipids, ROS, and calcium (Han *et al*., 2003; Bayrhuber *et al*., 2008; Reina *et al*., 2020). Our data indicates that alteration of VDAC homeostasis caused by tafazzin-deficiency disrupts calcium uptake into the mitochondria (Figure 4F&G), which is potentially causative for increased calcium levels in the cytosol (Figure 4F). ROS produced in mitochondria can no longer efficiently leave this compartment in a VDAC-dependent manner, which could explain their subsequent accumulation (Figure 1D & Figure 3C). Furthermore, tafazzin-deficient cells show a substantial reduction in ER-mitochondria contact sites (Figure 4H). The disruption of functional coupling between mitochondria and the remaining cell hinders mitochondria derived ROS to efficiently contributing to ferroptosis in tafazzin-deficient cells, leading to the observed protective effects (Figure 1F). Recently, Zhang et al. demonstrated that comparable ferroptosis suppressing effects can be triggered by pharmacological or genetic impairment of MAM formation via modulation of the IP_3_R-GRP75-VDAC1 complex (Zhang *et al*., 2024). In the case of tafazzin-deficiency, changes in cardiolipin metabolism and their effects on mitochondrial membrane properties cause comparable downstream disruptions.

While the trinity of reduced VDAC levels, their impaired oligomerization, and fewer ER-mitochondria contacts can be considered as the primary protective mechanism, the potential influence of other contributing factors needs to be considered. This includes the unsaturation of the membrane lipidome. Double bonds in fatty acyl residues are susceptible to (per-) oxidation by radicals generated from processes such as the Fenton reaction. Therefore, an increased content of PUFAs in membrane lipids accelerates ferroptosis (Qiu *et al*., 2024). Similarly, modulating enzymes that regulate double bond homeostasis in lipid metabolism, such as ACSL4 and LPCGAT3, have been shown to alter ferroptosis characteristics (Dixon *et al*., 2015; Doll *et al*., 2017). In accordance with this, we could modulate ferroptosis by altering the lipid environment, with linoleic acid having an accelerating and palmitic acid having a decelerating effect (Figure 3B). Tafazzin-deficiency reduces the double bond content in CLs but has no substantial impact on the abundance and the unsaturation index in other phospholipids (Figure 1B, Supplemental Figure 1&2) (Oemer *et al*., 2021), neither in mitochondrial-enriched nor microsomal fractions. Thus, the protective effect in the BTHS model cannot be explained via a phospholipid-dependent, ER-localized mechanism.

As an additional modulating factor, we considered alterations in iron metabolism, which is crucially involved in ferroptosis (Dixon *et al*., 2012). Mitochondria are key sites for iron storage and utilization (Lane *et al*., 2015). We investigated whether differences in cellular iron trafficking or storage could explain the protective effects (Figure 1F). However, in our model system tafazzin-deficiency did not alter intracellular and subcellular free iron availabilities, as assessed by the labile iron pool and total iron level measurements (Figure 2C, D&E), which are critical for determining the rates of the Fenton reaction. Furthermore, pharmacological modulation of iron levels did not mitigate the ferroptosis resistance observed in the BTHS model (Figure 2A).

In conclusion, we mechanistically explain how pathogenic alterations of mitochondrial membrane lipids due to erroneous CL remodeling are causing ferroptosis resilience in the presence of high oxidative stress levels. In order to contribute to ferroptosis, the mere formation of ROS and downstream lipid peroxides it not enough; they must also reach the correct subcellular compartments. Tafazzin-deficiency related mitochondrial membrane lipid alterations cause a functional impairment of VDACs, which are central players in the signaling between mitochondria and the remaining cell. Furthermore, VDACs are also crucial for the physical formation of contact sites between mitochondria and the ER. As a consequence, otherwise damaging molecules are trapped and accumulate in mitochondria, instead of driving ferroptosis. For overall cellular survival, this is a favorable situation, as mitochondria are evolutionarily adapted to handle elevated stress levels.

## Material and Methods

### Cell culture

If not stated otherwise, all cells were grown at 37°C, 100% humidity, and in an atmosphere of 5% CO_2_. HEK 293T (Ctrl) and ΔTAZ (ΔTAZ) were maintained in DMEM (1,000 mg/l D-glucose and sodium bicarbonate (D5546, Sigma Aldrich, St. Louis, USA) supplemented with 10% (v/v) heat-inactivated fetal calf serum (Fetal Calf Serum; 10500-064, Gibco, Invitrogen, Carlsbad, CA), 2 mM L-glutamine (17-605E, Lonza Group, Basel, Switzerland) and 100 U/ml Pen-Strep (17-602E, Lonza Group, Basel, Switzerland). To split and seed cells, the cell layer was washed with 1x DPBS (47302.03, Serva, Heidelberg, Germany), incubated with Trypsin/EDTA (T4174, Sigma Aldrich, St. Louis, USA) until the cells were fully detached, resuspended in medium and centrifuged for 5 minutes at 200 g. When cells were seeded for an experiment, cells were counted using the CellDrop 2 (DeNovix Inc., Wilmington, DE 19810 USA), trypan blue (T8154, Sigma Aldrich, St. Louis, USA) was used for live/death determination.

### Cardiolipin analysis with LC-MS/MS

Cardiolipin extraction and sample analysis were conducted as previously described in (Wohlfarter *et al*., 2022). As CL are lipids exclusive for mitochondria, no subcellular compartment enrichment strategies were used. Lipids were extracted following the Folch- method (Folch, Lees and Stanley, 1957) and extracted lipids were separated by RP-HPLC followed by time-of-flight mass spectrometric detection (TIMS-TOF-Pro, Bruker Daltonics, Bremen, Germany). The raw data was pre-processed and analyzed using a targeted method for feature integration in MZmine (Pluskal *et al*., 2010). Relative abundances were calculated using R (http://www.R-project.org) and visualized using GraphPad Prism 9 v9.0.1 (GraphPad Software, Inc) or R (Folch, Lees and Stanley, 1957).

### Phospholipid analysis with LC-MS/MS

Phospholipid sample analyses were performed as described in Koch et al., 2020 with modifications (Koch *et al*., 2020). Lipids contained in microsomal and mitochondrial enriched fractions were extracted by using the Folch method (Folch, Lees and Stanley, 1957). Lipid extracts from respective fractions were separated on an Elute HPLC system (Bruker Daltonics, Bremen, Germany) with the parameters summarized in Table 1 and were analyzed by time-of-flight mass spectrometric detection (TIMS-TOF-Pro, Bruker Daltonics, Bremen, Germany) with HESI (Vacuum Insulated Probe Heated Electrospray Ionization). More details about instrument setting are shown in Table 2. Bruker raw data files (.D) were converted by using an IBF converter into ion mobility base framework (IBF) format and then further processed in MS-DIAL (ver. 4.9.221218) as described in (Tsugawa *et al*., 2020). Data collection parameters were as follows: Retention time window: 4-18 min; Mass range (MS1 and MS/MS): 50-1300 Da, disregarding the RT parameters of the integrated library (Kind *et al*., 2013).

**Table 1:** Method details for Bruker HPLC Elute System and their respective settings for PL detection.

| HPLC system component | Settings |
| --- | --- |
| Column | Agilent Poroshell 120 EC-C8 2.7 mm 2.1x100 mm column (Agilent Technologies, Santa Clara, USA) |
| Column temperature | 50°C |
| Autosampler temperature | 10°C |
| Running solvent A | 10 mM ammonium formate and 0.2% formic acid in 6/4 (v/v) acetonitrile/H<sub>2</sub>O |
| Running solvent B | 10 mM ammonium formate and 0.2% formic acid in 9/1 (v/v) isopropanol/acetonitrile |
| Flow rate | 0.4 ml/min from 0 to 20 min 0.4-0.5 ml/ml for 2 min 0.5 ml/min for 3 min 0.4-0.5 ml/ml for 1 min 0.4 ml/min for 2 min |
| Gradient details | 40% B for 3 min 40-65% B for 17 min 65-99% B for 2 min 99% B for 3 min 99-40% B for 1 min 40% B for 2 min |
| Injection volume | 10 µl |

**Table 2:** Parameter details for mass spectrometric detection of PL and their applied settings.

| Parameters | Settings |
| --- | --- |
| Ionization | VIP-HESI in negative mode |
| Scan mode | MS/MS (PASEF) |
| Scan range | 100 m/z -1350 m/z |
| Nebulizer gas | 4.0 Bar |
| APCI Heater | 420 °C |
| End Plate Offset | -500 V |
| Dry Gas flow | 10.0 l/min |
| Dry Heater | 260 °C |
| Pre pulse storage | 5 μs |
| Collision Cell In | -70.0 V |
| Collision Cell Out | -25.5 V |
| Collision Cell RF | 1100.0 Vpp |
| Collision Energy (MS only) | -10.0 eV |
| IMS Ramp End | 1.57 V·s/cm<sup>2</sup> |
| IMS Ramp Start | 0.55 V·s/cm<sup>2</sup> |
| IMS Ramp Time | 100.0 ms |

Annotation of lipid species was carried out on basis of the following criteria: Reverse Dot product higher than 700 and Dot Product higher than 400 where MS/MS matched with integrated library (Kind *et al*., 2013). Then fragment spectra of each feature were manual cheeked to identify possible false positive annotations. Additionally, MS/MS peak list with m/z and relative intensity was exported from MSDial and the LipidMaps database search with a relative mass deviation of 5 ppm was carried out in MetFrag (Ruttkies *et al*., 2016).

Retention time (RT) correlation analysis was performed by linear regression between measured RTs in this study and the published RT values from Lange, et al., 2021 (Lange *et al*., 2021), as well as with an in house RT library (Koch *et al*., 2020). Features with an RT shift of >1 min were additionally checked if they fit in the lipid classes elution pattern. Confirmed phospholipids were normalize to 100% in a class-wise manner for each sample. Graphical representation was done in GraphPad Prism 10.2.2 (GraphPad Software, Inc).

### Life cell imaging

8000/well cells of Ctrl, tafazzin-deficient HEK cells (ΔTAZ) were seeded in a 96-well plate (Falcon, 353072) and incubated overnight. The following day, the cells were treated with 15 µM erastin (A13822, AdooQ Bioscience, Irvine, CA, USA) or 4 µM RSL3 (SML1414, Sigma-Aldrich, St. Louis, USA) with or without 4 µM Liproxstatin-1 (Lip-1) (SML1414, Sigma Aldrich, St. Louis, USA) with or without 20 µM Q-VD-Oph hydrate (QVD) (A14915, AdooQ Bioscience, Irvine, CA, USA). For live cell imaging with fatty acid treatment, the respective fatty acid was administered three days prior to the experiment, and the cells were seeded in the appropriate fatty acid supplemented medium and treated on the next day, as described. For Iron supplemented experiments 50 µM Fe (Iron(III) nitrate nonahydrate, Sigma F8508) and 50 µM DFP (Deferiprone, Sigma 379409-5g) were added simultaneously with erastin and the other compounds. The IncuCyte S3 microscope (Sartorius, Göttingen, Germany) with a 10x objective was used for live cell imaging, viability was determined using 1 µg/ml propidium iodide (PI) (81845, Sigma Aldrich, St. Louis, USA). During imaging, cells were kept in an incubator with a temperature of 37°C, 5% CO_2_ and 100% humidity, and images were captured at 2 h (erastin treated) or 30 min (RLS3 treated) intervals. Pictures were analyzed using the IncuCyte Software v2020B (Sartorius, Göttingen, Germany) and visualized using GraphPad Prism 9 v9.0.1 (GraphPad Software, Inc).

### Fatty acid supplementation

Fatty acid supplementation was carried out following the method described in (Oemer *et al*., 2020). Briefly, cells were supplemented with either BSA (vehicle control) (A7030, Sigma Aldrich, St. Louis, USA) conjugated with 25 µM palmitic acid (P0500, Sigma Aldrich, St. Louis, USA) or 25 µM linoleic acid (L1376, Sigma Aldrich, St. Louis, USA). To prepare a 100 mM fatty acid stock solution, the pure fatty acids were dissolved in 100 mM NaOH and the solution was incubated above the melting point of the fatty acid. This stock solution was then mixed with 2.8 mM BSA and diluted to 25 µM in cell culture medium before being added to the cells.

### Subcellular fractionation for AAS, aconitase assay & lipidomic profiling

The mitochondrial isolation procedure was conducted following the method described in (Wieckowski *et al*., 2009). The frozen cell pellets were homogenized by stroking with a needle tip 10 times in isolation buffer 1 (225 mM mannitol, 75 mM sucrose, 30 mM Tris-HCL pH 7.4, 0.5 mM EGTA) and centrifuged at 600 g for 5 minutes at 4°C. The supernatant was then transferred to a new vial and centrifuged again for 5 minutes at 600 g at 4°C. The resulting supernatant was collected as microsomal fraction (MSF). The remaining pellet was washed in isolation buffer 2 (225 mM mannitol, 75 mM sucrose, 30 mM Tris-HCL pH 7.4), centrifuged at 7000 g for 10 minutes at 4°C, and the supernatant was discarded. The pellet was then washed a second time in isolation buffer 2, again centrifuged at 10,000 g for 10 minutes at 4°C, and the supernatant was again discarded. The resulting pellet represents the mito-enriched fraction. Western blotting was used to ensure the purity of MSF and mito-enriched fraction.

### Mitochondrial isolation for respirometry and calcium uptake rates

For respirometry and other High-Resolution FluoRespirometry experiments (Oroboros Instruments, Innsbruck, Austria), mitochondria were isolated as described in (Frezza, Cipolat and Scorrano, 2007). Briefly, cells were collected, counted, and washed in PBS once. Centrifugation was carried out at 600 g for 10 minutes at 4°C. The supernatant was discarded, and the cells were suspended in 1.8 ml of ice-cold IB_c_ (10 ml of 0.1 M Tris/MOPS and 1 ml of 0.1 M EGTA/Tris to 20 ml of 1 M sucrose; volume was brought to 100 ml with distilled water, pH was adjusted to 7.4). Cells were then homogenized using a Teflon potter, with the cell suspension stroked 30 times. After another centrifugation step at 600 g for 10 minutes at 4°C, the supernatant was collected and transferred to a new 2 ml screw cap vial. This was followed by centrifugation at 7000 g for 10 minutes at 4°C. The supernatant was discarded, and the pellet was washed in 300 µl of ice-cold IB_c_. A further centrifugation at 7000 g for 10 minutes at 4°C was done, and the supernatant was discarded again. The mitochondrial pellet was resuspended in mitochondrial respiration medium MiR05 (Oroboros Instruments, Innsbruck, Austria) for O2k measurements.

### Calcium uptake capacity using HRR

Ca^2+^ uptake by isolated mitochondria was measured by fluorescence with 2 μM Calcium Green-5N (Thermo Fisher). The fluorescence was measured in 2-mL chambers of the NextGen-O2k fluorespirometer (Series XA, Oroboros Instruments) with a medium containing 110 mM sucrose, 70 mM KCl, 20 mM HEPES, 10 mM KH_2_PO_4_ , 1 mM MgCl_2_, pH 7.1 Smart Fluo-Sensors Blue were used with the MgG/CaG filter set (465 nm excitation, 250 intensity, gain 1000).

Prior to the addition of the samples, glutamate (10 mM), malate (2 mM), ADP (30 μM), and oligomycin (2.5 μM) were added to the chambers. EGTA was then titrated (10 μM per injection) to reduce the fluorescence signal. Once the fluorescence signal was not further reduced by EGTA, isolated mitochondria were reconstituted in IB_c_ and added to the chamber. CaCl_2_ was then titrated (0.5 μM per step). Upon CaCl_2_ titration, the fluorescence signal increases, followed by a decrease which reflects uptake of Ca^2+^ by mitochondria. CaCl_2_ was titrated until no more uptake was detected, and the fluorescence increased independently of the calcium addition, showing release of Ca^2+^ related to the mitochondrial permeability transition. The uncoupler CCCP (0.5 μM) was added at the end of the protocol to dissipate the proton motive force, if not yet complete by the mitochondrial permeability transition.

Ca^2+^ uptake was calculated by converting the fluorescence signal into Ca^2+^ concentration. The delta between the fluorescence signal immediately after CaCl_2_ injection and immediately before the following injection corresponds to the uptake. This fluorescence signal is converted into Ca^2+^ concentration by using the acute fluorescence signal increase caused by the previous CaCl_2_ injection of 2 nmol. The sum of each step corresponds to the total Ca^2+^ uptake, which was normalized by mitochondrial protein content (nmol Ca^2+^·mg^-1^).

### Cell fluorescence microscopy

For live cell analyses cells were grown on collagen-coated (0.1 mg/ml) glass slides. Images were collected using a 63x oil objective of an Axiovert200M microscope equipped with an ApoTome.2 system for structured illumination microscopy (Zeiss, Vienna, Austria) (Hagenbuchner and Ausserlechner, 2013; Hagenbuchner *et al*., 2016). Images were taken at equal exposure times. For quantification, at least 100 cells from three independent experiments were analyzed.

### Mitochondrial ROS quantification and morphology

For analyzing mitochondrial morphology, untreated, fatty acid-supplemented (or BSA control), or ferroptosis-induced cells were stained with CMXRos (M7512, 300 nM). Mitochondria with more than 50 branches per 30 × 30 µm were classified as "tubular," while mitochondria with more than 5 slubs/dots per 30 × 30 µm were classified as "fissioned." For ROS cells were incubated with CM-H_2_XRos (M7513, 500 nM; Invitrogen, USA).

Images were acquired with a Zeiss Axiovert200M microscope and cellular fluorescence intensity was quantified using Axiovision software (Zeiss, Vienna, Austria). Images were taken with equal exposure times and processed using Axiovision software.

### Lipid Peroxidation

For cellular lipid peroxidation levels, cells were incubated in 10 µM Bodipy-581/591 C11 (D3861, Thermofisher) for 10 minutes at 37°C, nuclei were stained with Hoechst (500 ng/ml).

### Cytosolic calcium

For cytosolic calcium measurements, cells were incubated with 2x concentrated Fluo-4 (see manufacturer’s manual for the Fluo-4 Direct™ Calcium Assay Kit F10472, Thermo Fisher Scientific) for 1 hour at 37°C and nuclei were stained with Hoechst (500 ng/ml).

#### Labile iron Pool imaging

For live cell analyses cells were grown on collagen-coated (0.1 mg/ml) slides as described above. Intracellular labile iron was detected using FerroOrange (Dojindo, Kumamoto, Japan) and for mitochondrial labile iron pool Mito-FerroGreen (Dojindo, Kumamoto, Japan) according to manufacturer’s protocol. Briefly, cells were washed twice with hanks, balanced salt solution (HBSS). Then cells were treated for 30 min at 37°C in the dark with 1 µM FerroOrange or 5 µM FerroGreen with HBSS. After incubation stains were removed by washing twice with HBSS. Images were detected using a Leica TCS SP8 confocal microscope, images were analyzed using the CellProfiler 4.2.6 software (Stirling *et al*., 2021).

### SDS-PAGE and immunoblotting

Harvested cells were lysed in RIPA buffer or the NP-40 Buffer (Thermo Fisher Scientific) (for VDAC crosslinking experiments). To do so, samples were incubated for 20 mins on ice with vortexing after 10mins. Debris was pelleted by centrifugation at 4°C at 13,000 g for 10 min and supernatant was transferred into a new vial. Protein was quantified using Bradford Protein assay (Biorad). Following the addition of LDS sample buffer, the samples were subjected to 70°C for 10 minutes. They were then loaded onto 4 -12% polyacrylamide bis-tris gel (Thermo Fisher Scientific) and transferred onto polyvinylidene difluoride membranes (Amersham) using the Thermo Fisher Scientific XCell II Blot module (Thermo Fisher Scientific). After 1h blocking at room temperature in 5% skim milk, the membranes were subsequently incubated overnight at 4°C in 5% skim milk with primary antibodies (GPx4 (Abcam, ab125066), VDAC1/3 (Abcam, ab14734), FPN (Eurogentec, self-made antibody), TfR1 (Invitrogen, 13-6800), PCBP1 (Abcam, ab74793), PCBP2 (Antikörper-online, ABIN3023340), IRP2 (Novus Biologicals, NB100-1798), MnSOD1 (Enzo Life Science, ADI-SOD-110), ABCB7 (Novus Biologicals, NBP2-21600), Actin (Sigma, A 2066), Mitoferrin2 (BiossAntibodies, bs-7157R), α-Tubulin (Abcam, ab7291), GAPDH (Cell Signalling Technology, #2118), followed by incubation at room temperature for 1 hour with secondary antibodies coupled to horseradish peroxidase (HRP). The HRP-conjugated secondary antibodies were visualized using Clarity Western ECL (Bio-Rad). Immunoblots were captured using Amersham equipment. Western blot bands were quantified using Fiji (Schindelin *et al*., 2012).

### Chemical VDAC crosslinking

Cells were exposed to 15 µM of erastin for 0h or 8h and then harvested without using trypsin. Subsequently, they were washed twice with PBS and reconstituted in 200 µM EGS (21565, ThermoFisher Scientific) solution in PBS (pH=8.3) and incubated at 30°C for 15 minutes. Afterwards, cells were pelleted through centrifugation at 10,000 g for 5 minutes. 50 µg of protein was further subjected to SDS-Page and immunoblotting, employing the anti- VDAC1/Porin +VDAC2 antibody (ab154856) (Baik *et al*., 2023).

### High-resolution respirometry

Cells were treated and cultivated in 25 cm^2^ flasks, counted and mitochondria were isolated as described above. Two instruments were used in parallel and each instrument is composed of two 2 ml chambers thermostatted at 37°C. Respiration medium was equilibrated with air for calibration of the oxygen sensors. The oxygen range was maintained between air saturation and a minimum of 100 µM Oxygen flux [pmol O2^·s-1^·mL^-1^] was automatically corrected for instrumental background and results were expressed as oxygen flow per cell [amol O2·^s-1^·cell^-1^]. A substrate-uncoupler-inhibitor titration protocol (SUIT-008), which was designed to assess the additivity between the N- and S-linked pathways in the Q-junction to provide a physiologically relevant estimate of the maximum mitochondrial respiratory capacity. Briefly, mitochondria were added in MirO5 (1mt). Then 5 mM pyruvate and 2 mM malate (PM) to support N-linked was added, yet in the absence of ADP, ATP and AMP resulting in a non-phosphorylating resting state (LEAK state). The addition of 2.5 mM ADP induces active N-OXPHOS capacity (ADP). 10 µM cytochrome c (c) was added to test for the loss of integrity of the OMM, which would be indicated by a stimulation of respiration. 10 mM L-glutamine (G) was added to increase N-linked OXPHOS capacity. Addition of 10 mM succinate (S) stimulates the simultaneous NS-OXPHOS capacity with convergent electron flow from both, RCI and RCII in the Q-junction. The uncoupler CCCP (0.5 - 1 µM carbonyl cyanide m-chloro phenyl hydrazine) is titrated stepwise until the maximum level of electron transfer (ET) capacity has been reached entering the ET-state. 1 µM rotenone inhibits CI and eliminates N-linked substrates as electron donors resulting in a solely S-linked control state (Doerrier *et al*., 2018).

### Enzyme activity assays

Aconitase activity was measured in isolated mitochondria (described above) using the Abcam Aconitase Activity Assay Kit (ab109712) according to manufacturers instructions. 60 µg mitochondria were assayed in a 96-well UV plate provided and incubated with isocitrate and manganese. The conversion of isocitrate to cis-aconitate was recorded by a kinetic measurement at OD_240nm_ for 30 minutes. The resulting aconitase activity was calculated as recommended in the manual.

Citrate Synthase activity was measured as described in (Brischigliaro *et al*., 2022). The Citrate Synthase assay was conducted using a reaction mix composed of 100 mM Tris-HCl + 0.2% Triton X (pH=8), 150 µM DTNB, 300 µM Acetyl-CoA, and 500 µM Oxaloacetate. Mitochondria were diluted to a final concentration of 1 µg/µl and added to the reaction mix, excluding Oxaloacetate. The mixture was incubated at 30°C for 5-10 minutes before being transferred to a 96-well plate. Baseline measurements were taken at OD_412nm_, followed by the addition of Oxaloacetate. Absorption was measured for at least 2 minutes at OD_412nm_. Data analysis involved path length correction and selection of a linear area, with calculations based on the formula (ε=13.8mM^-1^cm^-1^):

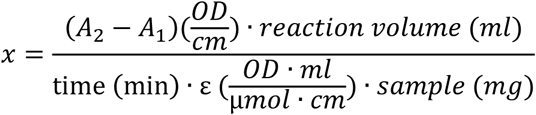

### Iron atomic absorption spectroscopy

Intracellular iron content was quantified using atomic absorption spectroscopy (AAS), following the method established by (Gehrer *et al*., 2022; Bernkop-Schnürch *et al*., 2023). Briefly, the cell pellets were resuspended in Milli-Q water, 0.2% Triton X-100 and lysed by sonication in a cup booster (Sonopuls, Bandelin) with cooling at 4 °C. The intracellular iron content was quantified by graphite furnace atomic absorption spectrometry (M6 Zeeman GF95Z-AA-Spectrometer; Thermo Fisher Scientific) using Extended Lifetime graphite cuvettes. Measurements were done at 248.3 nm (bandpass 0.2 nm) under argon atmosphere using 1100°C ash temperature and 2100°C atomization temperature with Zeeman background correction.

### Mitochondria-ER contact sites

To monitor ER-mitochondria interactions and morphologies, cells were seeded at 60% confluency into 6-well plates. After letting the cells adhere to the surface for at least 12 hours, cells were transfected with SPLICS Mt-ER Short P2A plasmid DNA (Addgene, #164108) using Mirus Bio™ TransIT™-LT1 Transfection Reagent. After 24h, cells were incubated with 200nM MitoTracker Deep Red FM (Invitrogen) for 30 min at 37°C prior to fixation. The cells were then fixed with 4% PFA for 20 min at RT, washed twice with 1x PBS, permeabilized with 0.1% Triton X-100 for 3 min, washed twice with 1x PBS, blocked with 2% BSA, and incubated with anti-Calnexin primary antibody (Abcam) for 1h at RT. After primary antibody incubation, cells were washed twice with 1x PBS and then incubated with goat anti-rabbit Alexa 568 conjugated secondary antibody (Sigma) for 1 hour at RT. Lastly, samples were washed with PBS and mounted using polyvinylalcohol (Sigma). Imaging was performed on a Nikon Crest-V3 spinning disc confocal microscope using a 60x objective (CFI Apochromat TIRF 60× NA: 1.49). A complete z-stack of the cell was acquired every 0.1 µm. Z-stacks were processed using Fiji (Schindelin *et al*., 2012). Images were filtered and processed using functions of the 3D image suite library (Ollion *et al*., 2013). For each condition, a minimum of 50 cells were quantified.

### Glutathione measurements by LC-MS/MS

#### Sample preparation for GSH LC-MS/MS measurements

Cells were exposed to 15 µM of erastin, with and without additional 4 µM Lip-4 and 20 µM QVD, and pelleted after 0h, 5h, 24h, 29h and 48h. Following cell harvest, samples were treated with 2 mM of the glutathione (GSH) masking agent N-ethylmaleimide (NEM) for 5 minutes to prevent the autooxidation of GSH to glutathione-disulfide (GSSG). Subsequently, cells were pelleted by centrifugation at 1000 g for 5 minutes, and the supernatant was removed before snap freezing the samples in liquid nitrogen. For LC-MS/MS extraction cells were reconstituted in 300 µl MeOH containing 0.5 µM GSH standard (Cambridge Isotope Laboratories, Inc., Andover, USA), and glass beads were added for cell lysis by shaking at 20Hz for 2.5 minutes. Further homogenization was achieved by a 5-minute ultrasonication step. To remove cell debris samples were centrifuged at 4°C, 31150 g for 10 min. The resulting supernatant was then transferred into a new glass vial, and organic solvents were evaporated using a nitrogen flow, followed by storage at -80°C. Prior to LC-MS/MS measurements, samples were reconstituted in H_2_O.

#### GSH LC-MS-MS method

Experiments were performed on a trapped ion mobility spectrometry (tIMS) time-of-flght (TOF) mass spectrometer coupled to a Bruker Elute uHPLC (Bruker Daltonics, Bremen Germany) combined with an Atlantis Premier BEH C18/AX 1.7 µm, column (Waters Corporation, Milford, USA), following LC and MS parameters were used (Table 3):

**Table 3:**
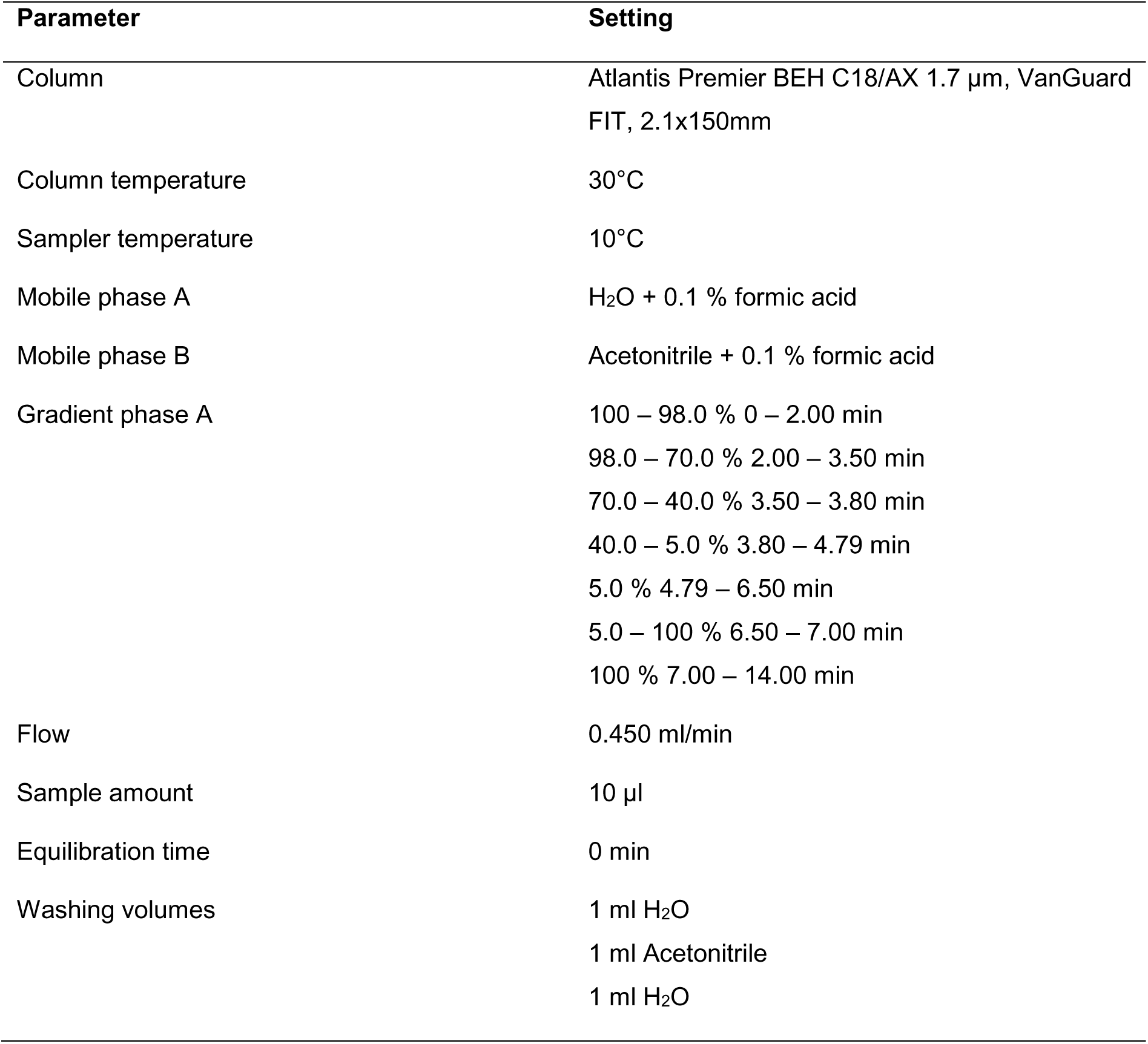
Method details for Bruker HPLC Elute System and their respective settings for GSH detection.

For the MS settings (Table 4), metabolomics Bruker optimized settings were adopted, and the mass range was adjusted. For mass calibration, a quadratic calibration with a tuning mix was used.

**Table 4:** Parameter details for mass spectrometric detection of GSH and their applied settings.

| Parameters | Settings |
| --- | --- |
| Ionization | Apollo II ESI in positive mode |
| Scan mode | bbCID(MS-MS/MS) |
| Scan range | 50 – 800 m/z |
| End plate offset | 500 V |
| Capillary | 4500 V |
| Nebulizer | 2.5 bar |
| Dry gas | 10.0 l/min |
| Dry temp. | 200°C |
| Deflection 1 delta | 70.0 V |
| Funnel 1 RF | 350 Vpp |
| Funnel 2 RF | 600 vpp |
| Multiple RF | 400 Vpp |
| isCID energy | 0.0 eV |
| Ion energy | 5.0 eV |
| Low mass | 200 m/z |
| Collision energy | 7.0 eV |
| Collision RF | 1500 Vpp |
| Transfer time | 100 µs |
| Pre pulse storage | 5 µs |

Targeted GSH data was analyzed using TASQ (Bruker Daltonics). Absolute quantification was performed using an external calibration curve with seven different concentrations ranging from 0.00305 to 12.5 µM. The obtained values were then normalized to protein content.

### Modeling of the spatial distribution of ferroptosis in live cell imaging experiments

#### Data processing

Brightfield and fluorescence channel images from the IncuCyte experiments were exported separately and processed using scipy (Virtanen *et al*., 2020) and scikit-image (van der Walt *et al*., 2014). The brightfield data was segmented into foreground (entropy > 5) and background (entropy ≤ 5) regions using pixel entropy. Foreground regions of interest (ROIs) were expanded by 25 μm. These expanded regions were labeled and mapped back onto the initial foreground. Dead cell signals from the fluorescence channel were identified using the difference of Gaussians (DoG) algorithm, with a threshold of 2.5 and a maximum σ of 5.

### Experimental radial distribution calculation

The experimental radial distribution of the PI+ cells was calculated by determining all pairwise distances between dead cells within each ROI. These distances were then used to construct a histogram representing the radial distribution.

### Modeling cell death

Ferroptotic cell death was modeled by iteratively selecting ROI pixels as dead cells. To obtain a robust histogram efficiently, this random sampling was repeated 100 times. The simulated histograms were constructed from the randomly drawn “dead cells” similar to the experimental radial distribution.

### Positive control model

The positive control model simulates a higher probability of cell death near other dead cells. Therefore, the probability of drawing a ROI pixel changes during the simulation based on its distance to other drawn pixels. After an initial draw, pixels within a radius *r*_*i*_ of the drawn pixel cannot be drawn again, mimicking dead cell size. Pixels within *r*_*i*_ and *r*_*o*_ have an increased (*f* times) probability to be drawn, while the probability of pixels beyond *r*_*o*_ remains unaltered. Probabilities are normalized after adjustment.

### Negative control model

The negative control model simulates unbiased cell death. While this could be calculated analogously to the positive control model, a computationally faster approach is used. For each ROI, the radial distribution of its pixel is calculated using the scikit-learn (Pedregosa *et al*., 2011) KDTree implementation. The obtained radial distribution is scaled by the number of pairwise distances between the detected dead cells in the ROI. The scaled radial distribution is summed up across all ROIs to obtain the histogram representing the underlying probability distribution.

### Supercomplex analysis

Complex and supercomplex levels, compositions, and distributions were analyzed in isolated mitochondria by blue native polyacrylamide gel electrophoresis (BN-PAGE) (Acín-Pérez, Hernansanz-Agustín and Enríquez, 2020). Briefly, isolated mitochondria were solubilized with 10% digitonin (4 g/g; Sigma-Aldrich D5628) and resolved in 3-13%-gradient Blue native gradient gels (1.5-mm thick), which were prepared using a gradient former connected to a peristaltic pump.

BN-PAGE proteins were electroblotted using a Mini Trans-Blot Cell system (Bio-Rad) onto polyvinylidene difluoride (PVDF) transfer membrane (Immobilon-FL, 0.45 μm; Merck Millipore, IPFL00010) for 1 hour at 100 V in transfer buffer (48 mM Tris, 39 mM glycine, and 20% ethanol). Non-specific binding sites were blocked by incubating membranes with phosphate-buffered saline (PBS) containing 5% bovine serum albumin (BSA) for 1h at RT. For protein detection, membranes were incubated in blocking buffer containing primary antibody overnight at 4°C. After three washes with PBS 0.1 % Tween-20 (PBS-T) for 30 min, membranes were incubated with appropriate secondary antibodies for 1 hour at RT. Membranes were then washed twice with PBS-T and once with PBS. To study complex and supercomplex assembly, the PVDF membrane was sequentially probed with antibodies against CI (anti-NDUFA9, Abcam ab14713), CIII (anti-UQCR2, Proteintech, 14742-1 AP) and CIV (anti-COI, Invitrogen, MTCO1 459600). Antibody binding was detected by fluorescence as previously described (Acín-Pérez, Hernansanz-Agustín and Enríquez, 2020).

## Acknowledgements

We thank Vanessa Rechner for her outstanding technical assistance, Katrin Watschinger for her valuable insights and support regarding BH_4_ experiments and Andreas Villunger for the possibility to use the IncuCyte infrastructure.

## Contributions

Y.W. and M.A.K. conceived and designed the study and wrote the manuscript. Y.W., J.H., U.H., G.Ö., A.W., M.S., J.S., P.H.A., V.J., L.F.G.S., and H.T. performed experiments. Y.W., J.H., U.H., G.Ö., A.W., M.S., J.S., J.K. P.H.A., V.J., L.F.G.S., and H.T. analyzed experimental data. Y.W., J.H., U.H., H.F., P.H.A., J.A.E.D., T.E.A., G.W., and M.A.K interpreted data. Y.W., J.H., U.H., P.H.A., J.A.E.D, H.F., T.E.A., G.W., J.Z., and M.A.K. provided resources. All authors revised and agreed on the content of the paper.

