## Supplementary Material for "Mitochondrial cardiolipin metabolism controlled by tafazzin enables ferroptosis"

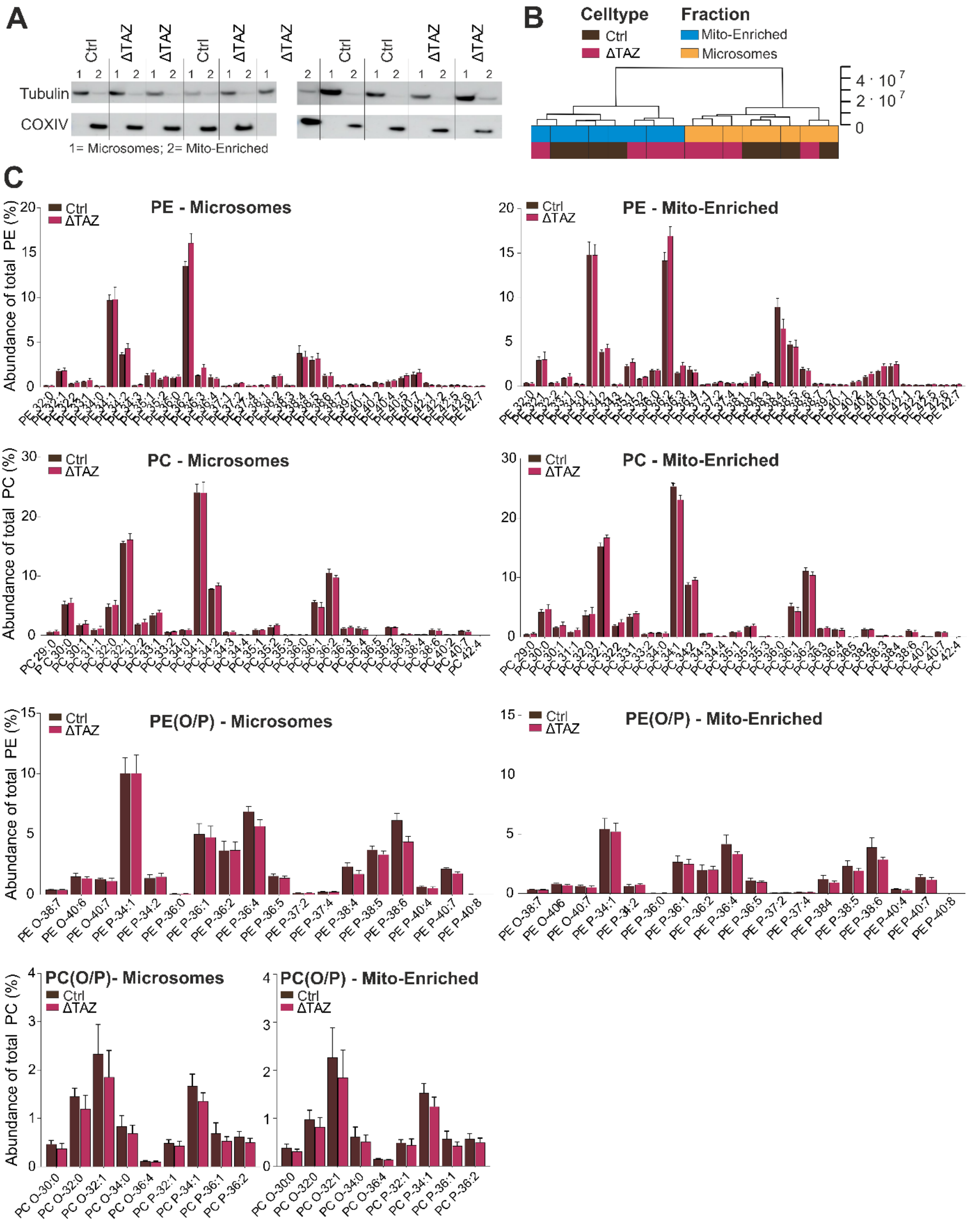


**Supplementary Figure 1: Phospholipidomic profiling in subcellular fractions of Ctrl and ΔTAZ cells.** A) Western blot analysis of subcellular fractionation efficiency using COXIV as mitochondrial and tubulin as microsomal marker. B) Dendrogram representing subcellular fractionation of Ctrl and ΔTAZ cells where Euclidean was used to measure distance and Ward as a clustering algorithm. C) Relative Phospholipid abundances in mito-enriched in microsomal fractions (%) normalized to individual phospholipid class. Phosphatidylcholines (PC), ether-linked PC (PC(O/P)) phosphatidylethanolamines

(PE), ether-linked PE (PE(O/P)). PE bar charts represent the 35 most abundant species. Data represented as mean±sd (n=4).

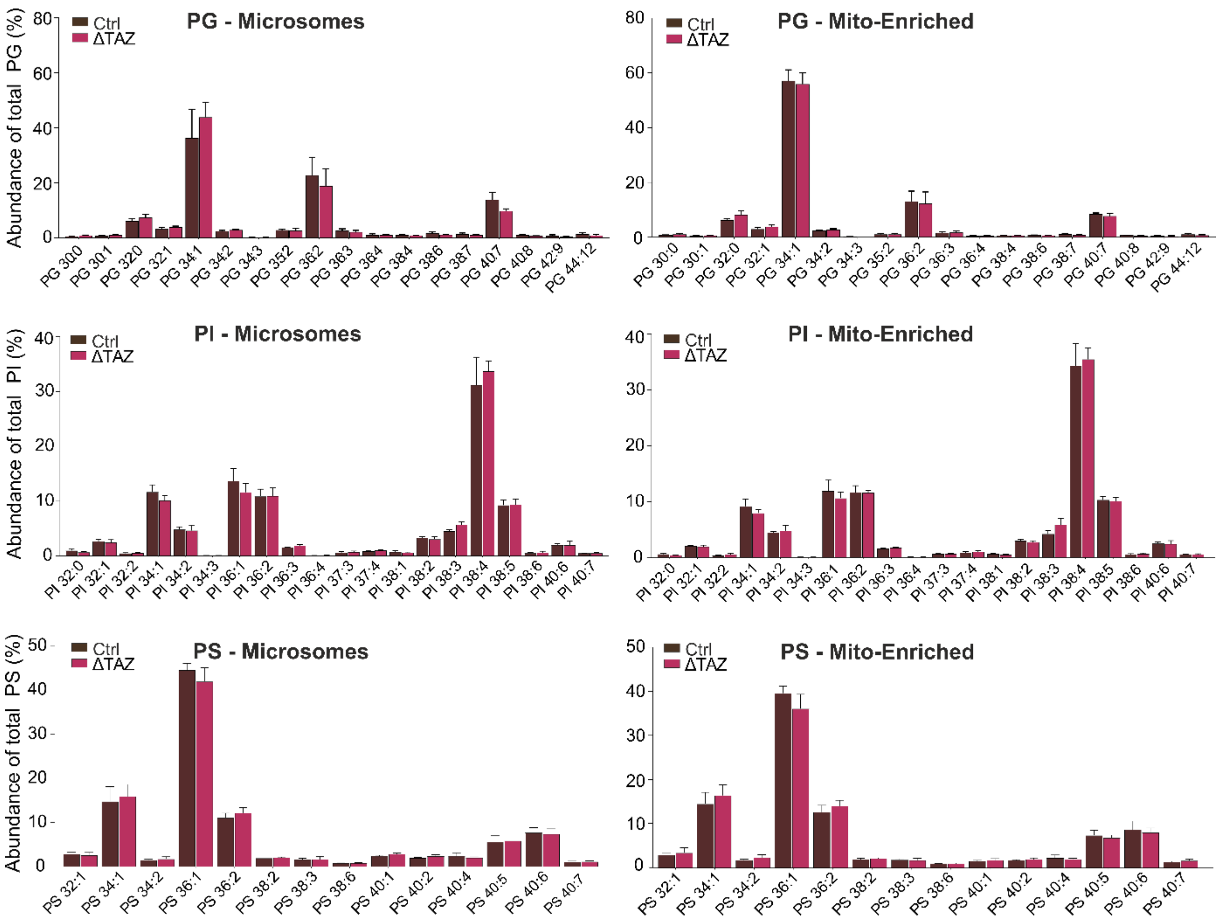


**Supplementary Figure 2: Phospholipidomic profiling in subcellular fractions of Ctrl and ΔTAZ cells**. Relative Phospholipid abundances in mito-enriched in microsomal fractions (%) normalized to individual phospholipid class, phosphatidylserines (PS), phosphatidylglycerols (PG), phosphatidylinositols (PI). Data represented as mean±sd (n=4).

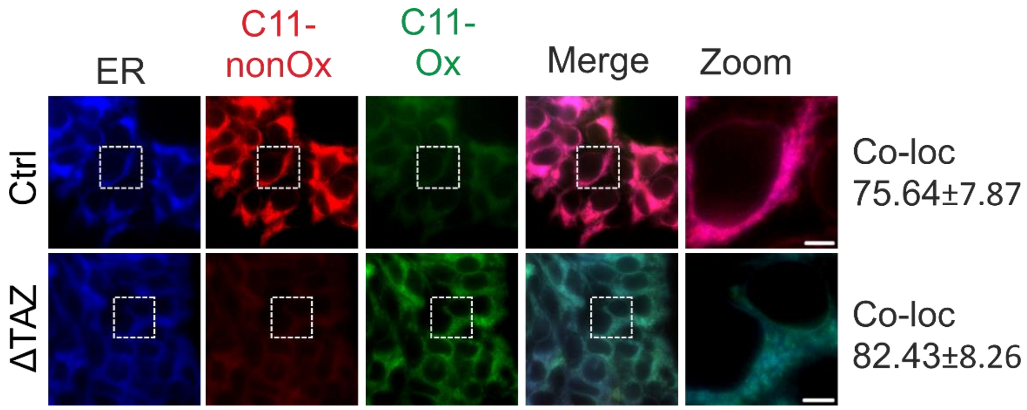


**Supplementary Figure 3: Cell imaging of Ctrl and ∆TAZ cells with Bodipy 581/591 C11 and ER staining.** Cell imaging of HEK and HEK∆TAZ cells was performed using Bodipy581/591 C11 to assess lipid peroxidation. The ER was stained with 333 nM for 10 minutes at 37°C using ER-TrackerTM Blue-White DPX (E12353, ThermoFisher Scientific). Co-localization of Bodipy 581/591 C11 and ER staining was calculated using AxioVision software (Zeiss, Vienna, Austria). Scale bar represents 5 µm.

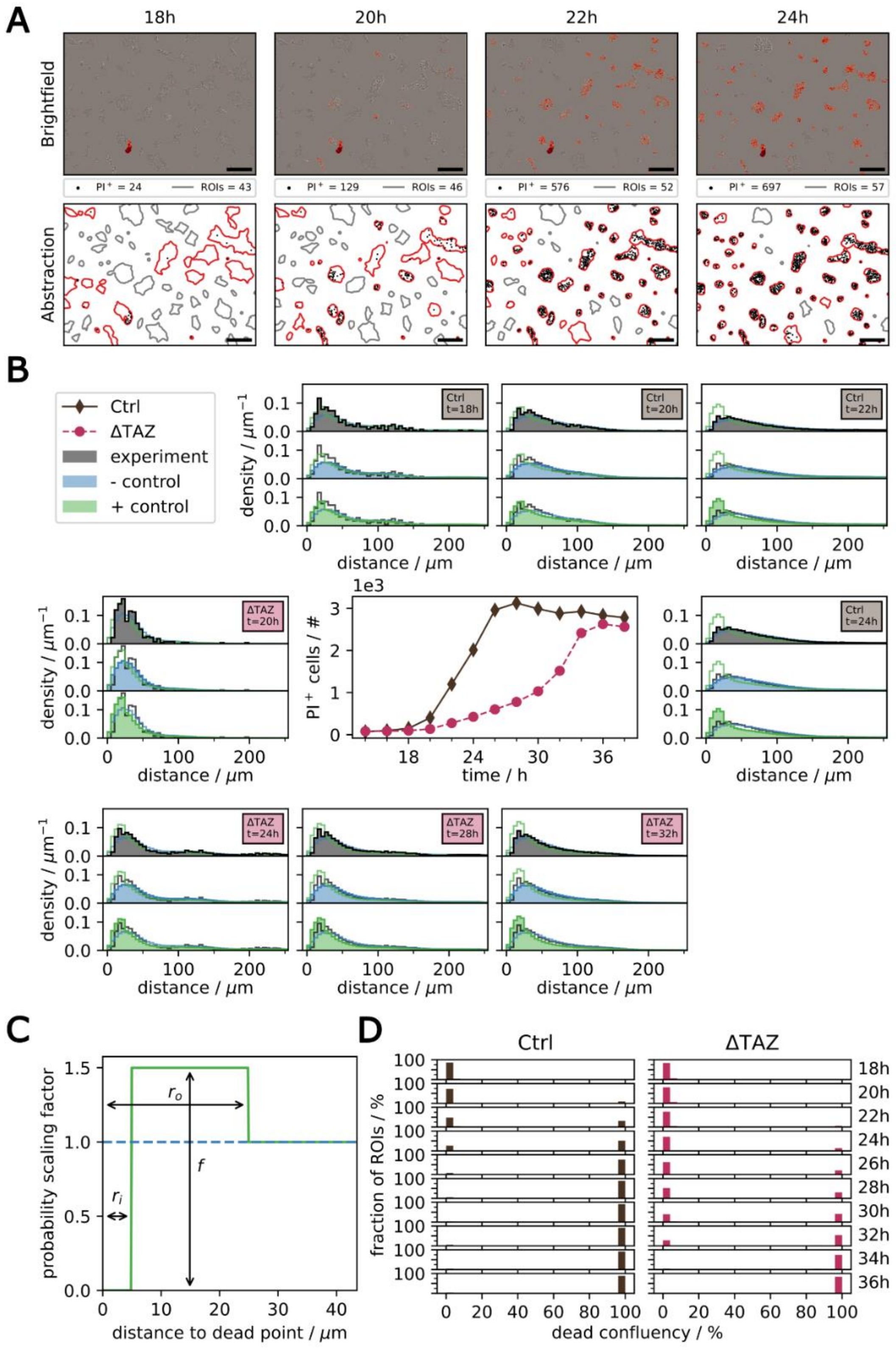


**Supplementary Figure 4: Modelling of the spatial distribution of ferroptosis in Incucyte experiments in Ctrl and ΔTAZ.** A) Abstraction of the bright field microscopy image and PI+ signal (upper part, bar size 200 µm) into regions of interest (ROIs) and identified dead cells. Foreground segments that are within 50 µm of each other are grouped into one ROI. B) Normalized radial distribution of identified dead cells within their respective ROI for Ctrl and ΔTAZ calculated from experimental data (in black), a negative (in blue), and a positive computational model (in green). The positive model parameters are f=1.5, $r_i$=2.5 µm, and $r_o$=25 µm. The chosen time points for Ctrl and ΔTAZ contain a similar number of $PI^+$ cells. C) Change in probability of surrounding pixels to be drawn in the next simulation step as a function of distance of the random start point for the negative (blue) and positive control (green) model. D) Histogram of the experimental dead cell confluency per ROI.

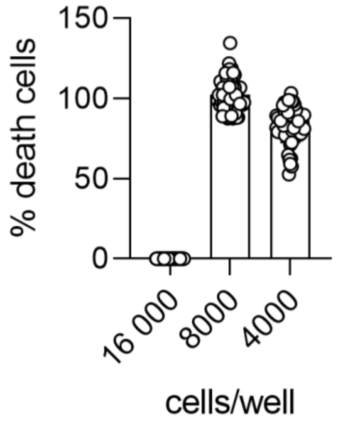


**Supplementary Figure 5: Cell death induction in Ctrl cells with 15 µM erastin**. Cell death induction in Ctrl cells treated with 15 µM erastin was measured using live cell imaging at 72 hours. The assay was conducted under three different seeding densities: 16,000, 8,000, and 4,000 cells per well in a 96-well plate. Cell death was indicated by the presence of propidium iodide (PI+) at a concentration of 1 µg/ml. The results showed that at the highest seeding density of 16,000 cells per well, the cells were too dense to undergo significant cell death. In contrast, at lower seeding densities of 8,000 and 4,000 cells per well, a higher degree of cell death was observed. Data points represent the mean ± sd.

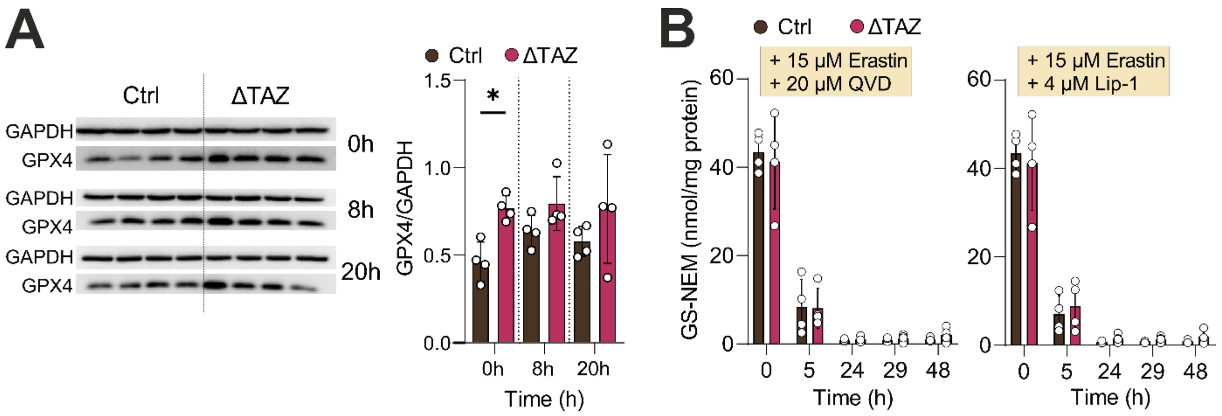


**Supplementary Figure 6: Ferroptosis induction control experiments. A)** GPx4 levels quantified via western blotting at 0, 8, and 20 hours after 15 µM erastin treatment (Ordinary one-way ANOVA, multiple testing corrected, p=0.044). **B)** GS-NEM depletion over 48h after 15µM erastin ± QVD ± Lip-1 treatment in Ctrl and ΔTAZ cells was measured via LC-MS/MS in 4 different tafazzin knock outs and 4 different controls, no significant differences could be observed between controls and knock out cells.

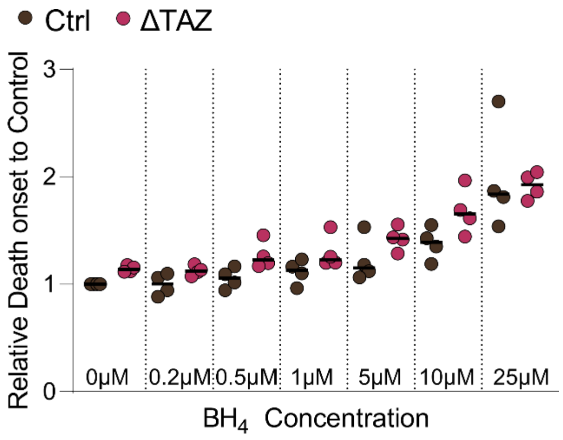


**Supplementary Figure 7: Effect of $BH_4$ on 15 µM erastin-induced cell death.** Cell death induction in Ctrl and ΔTAZ cells treated with 15 µM erastin was measured using live cell imaging. The relative onset of cell death compared to control was assessed in the presence of varying concentrations of BH4: 0 µM, 0.2 µM, 0.5 µM, 1 µM, 5 µM, 10 µM, and 25 µM. Data indicate that increasing concentrations of BH4 modulate the timing of cell death onset in Ctrl ΔTAZ cells treated with erastin. The delayed ferroptosis onset of ΔTAZ cells compared to controls remained throughout the experiment.

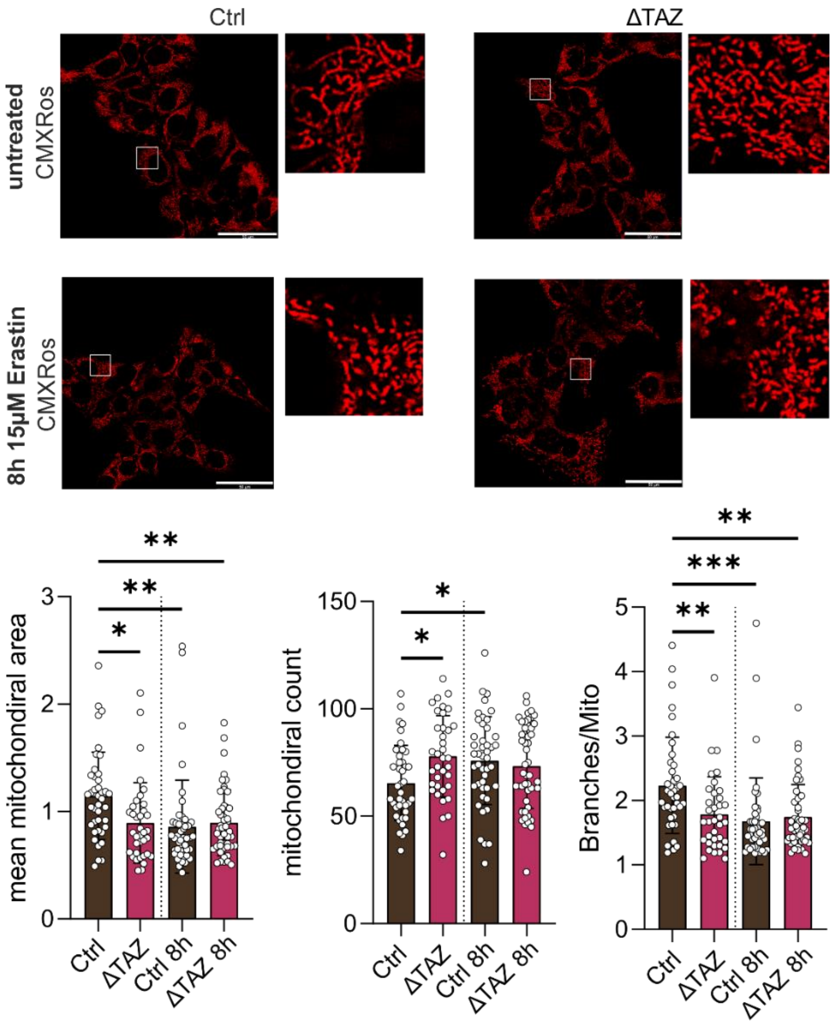


**Supplementary Figure 8: Mitochondrial fragmentation during ferroptosis.** Mitochondrial morphology was stained with CMXRos Mitotracker Red (200 nM) and analyzed using the "Mitochondrial Analyzer" in Fiji (1). In ΔTAZ cells, mitochondria were fragmented initially and remained fragmented following erastin treatment. In contrast, Ctrl cells displayed no mitochondrial fragmentation at the beginning, but fragmentation was induced after the treatment, reaching a level comparable to that observed in ΔTAZ cells. (Ordinary one-way ANOVA, multiple test corrected, $p<0.001$(***), $p<0.002$(**), $p<0.033$(*).

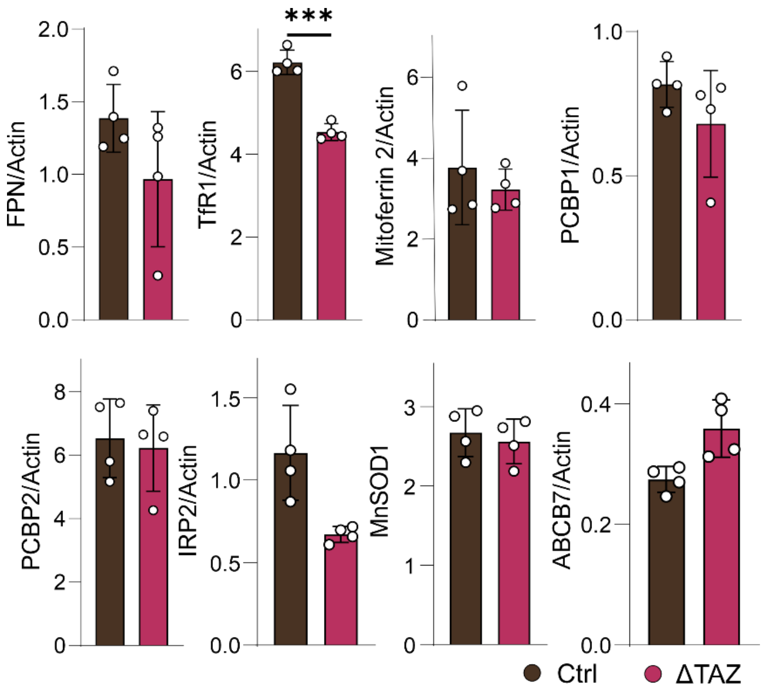


**Supplementary Figure 9: Iron transporter expression levels in Ctrl and ΔTAZ cells.** Levels of ATP binding cassette subfamily B member 7 (ABCB7), Ferroportin (FPN), Transferrin receptor 1 (TfR1), Iron regulatory protein 2 (IRP2), mitochondrial superoxide dismutase (MnSOD), and Poly C Binding Protein 1/2 (PCBP1/PCBP2) were measured via western blotting in Ctrl and ΔTAZ cells, across four different tafazzin knockouts and four different controls and quantified in ImageJ. Among these proteins, only the transferrin receptor (TfR1) was significantly lower expressed in ΔTAZ cells), (Two-way ANOVA, multiple testing corrected, $p<0.001$).

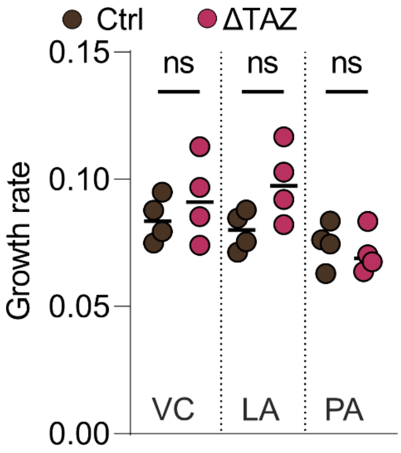


**Supplementary Figure 10: Growth rates of Ctrl and ΔTAZ cells remained constant upon lipid supplementation.** Growth rates of Ctrl and ΔTAZ cells in lipid were evaluated in the presence of 25 µM linoleic acid (LA), 25 µM palmitic acid (PA), or vehicle control (VC). Growth was assessed using the Logistic growth model in GraphPad Prism 9. No significant differences in h were observed between Ctrl and ΔTAZ cells under these conditions (Ordinary one-way ANOVA, multiple test corrected).

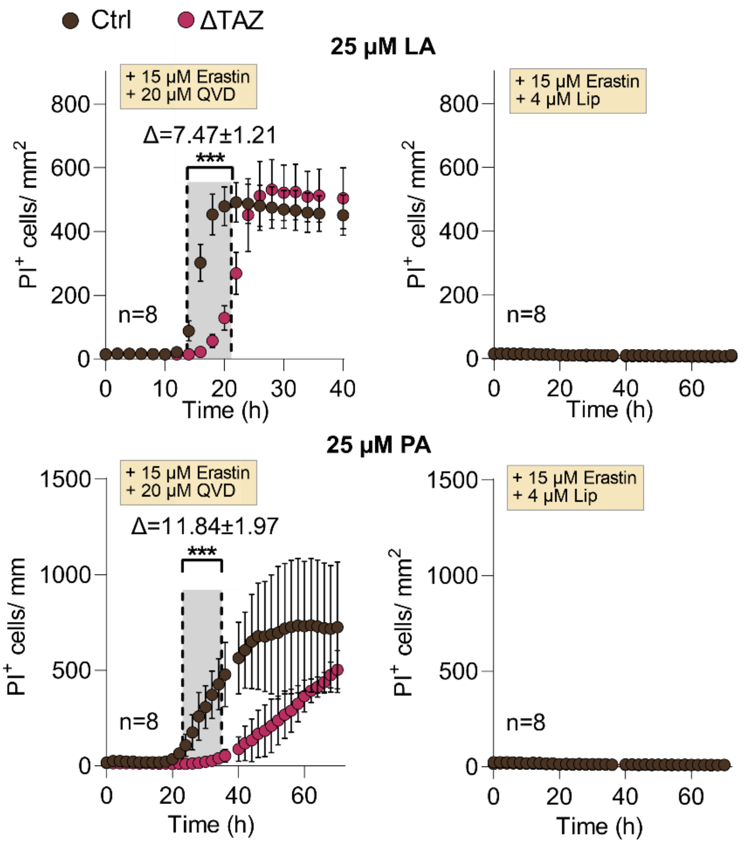


**Supplementary Figure 11:** Cell death induction in Ctrl and ΔTAZ with 15 µM erastin ± 20 µM QVD ± 4 µM Lip-1 in combination with 25 µM linoleic acid (LA) or 25 µM palmitic acid (PA) followed by live cell imaging over 72h, cell death was indicated via $PI^+$ (1 µg/ml), (Two-way ANOVA, multiple testing corrected, $p<0.001$).

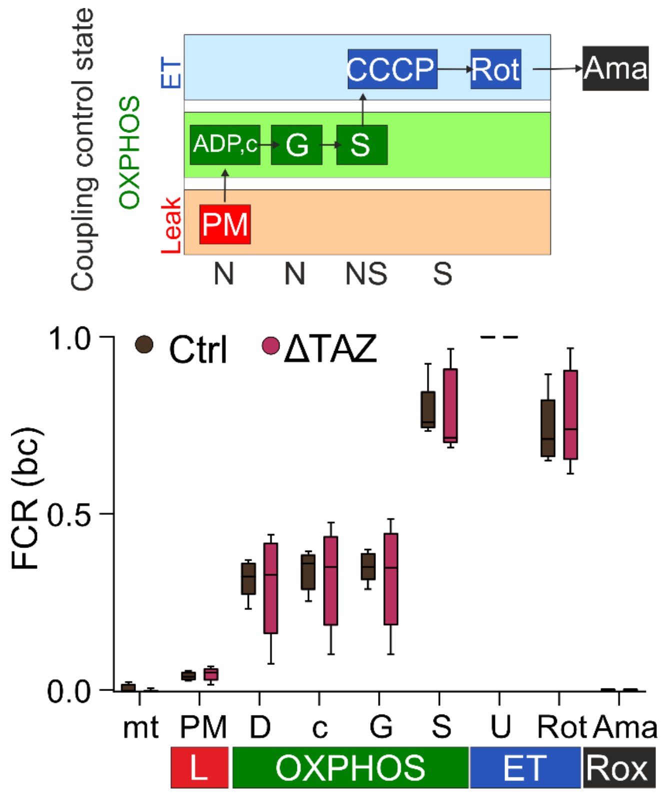


**Supplementary Figure 12: High-resolution respirometry** (O2K-FluoOxygraph, Oroboros Instruments) in isolated mitochondria with substrate-uncoupler-inhibitor titrations (SUIT-008 protocol). FCR = flux control ratio (baseline corrected with residual oxygen consumption after adding PM = addition of pyruvate and malate as substrates (NAD(P)H-dependent OXPHOS), D = addition of ADP for OXPHOS activation, c = addition of cytochrome c to control intact mitochondrial membranes, G = addition of glutamate as substrate (NAD(P)H-dependent OXPHOS), S = addition of succinate (NAD(P)H-dependent & succinate-dependent OXPHOS), U = addition of CCCP for membrane uncoupling (full electron transfer capacity, normalized to 1). R = addition of rotenone (CI inhibitor, for determination of succinate-dependent respirometry) and Antimycin A (Ama, CIII inhibitor), Shown is the mean±sd n=5.

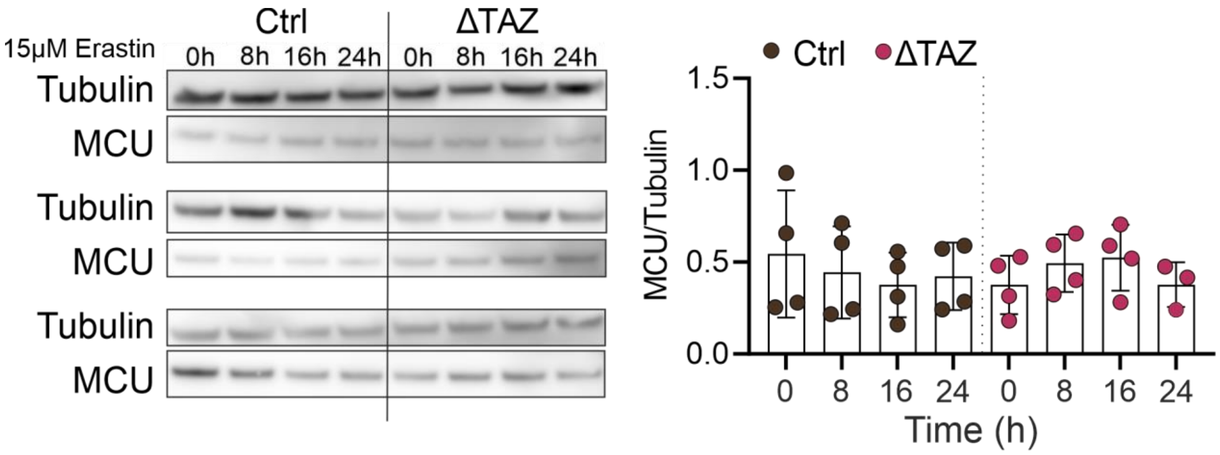


**Supplementary Figure 13: Mitochondrial calcium uniporter (MCU) levels** in Ctrl and ΔTAZ cells after 0, 8, 16, 24h of 15 µM erastin treatment. Throughout this time course, no upregulation of MCU was observed in response to ferroptosis induction in either cell type. This contrasts with the behaviour of VDAC1/3, which showed an upregulation following erastin treatment. The lack of MCU upregulation suggests that, unlike VDAC1/3, MCU does not play a role in the cellular response to ferroptosis induced by erastin in these cells.

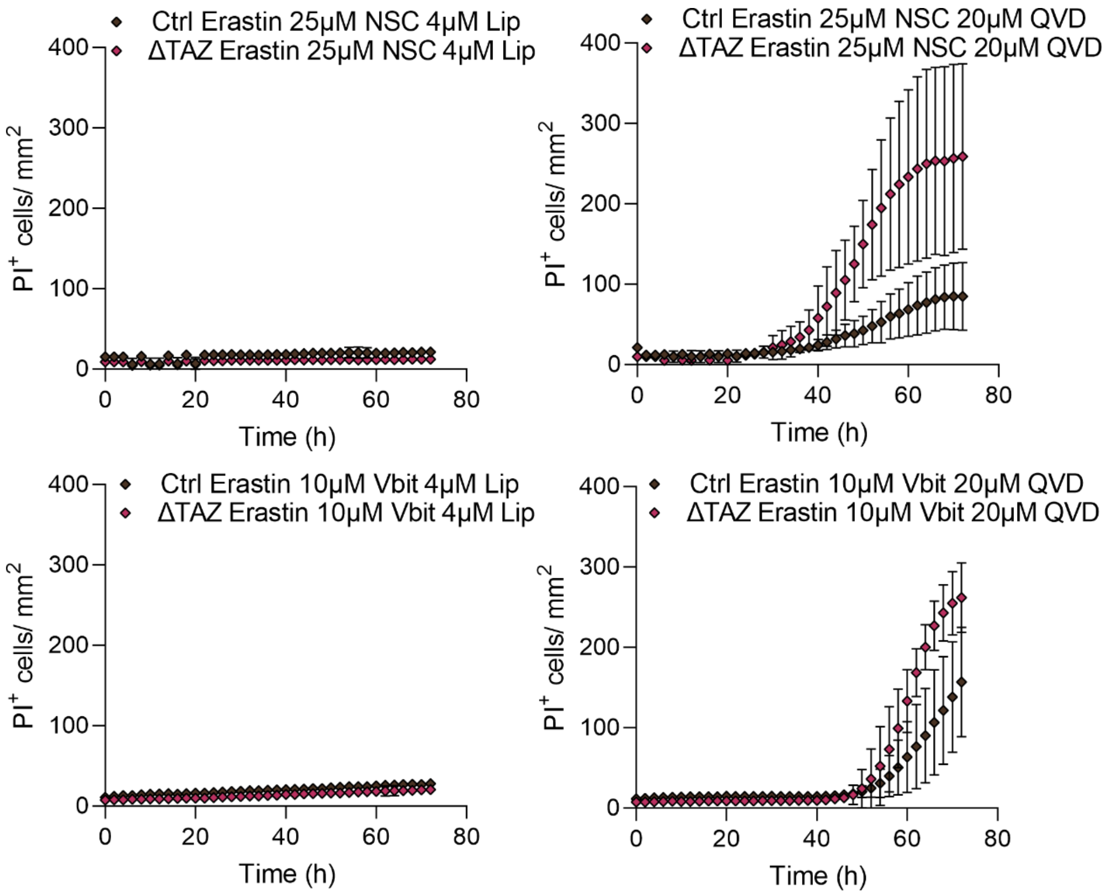


**Supplementary Figure 14:** Cell death was induced in Ctrl and ΔTAZ cells using 15 µM erastin, with additional treatments of 20 µM QVD, 4 µM Lip-1, 25 µM NSC, 10 µM Vbit4. Live cell imaging was conducted over 72 hours, with cell death indicated by $PI^+$ (1 µg/ml) staining. Lip-1 fully inhibited cell death, while cells additionally treated with QVD still underwent ferroptosis. These results indicate that Vbit4 and NSC do not alter the form of regulated cell death but specifically delay ferroptosis.

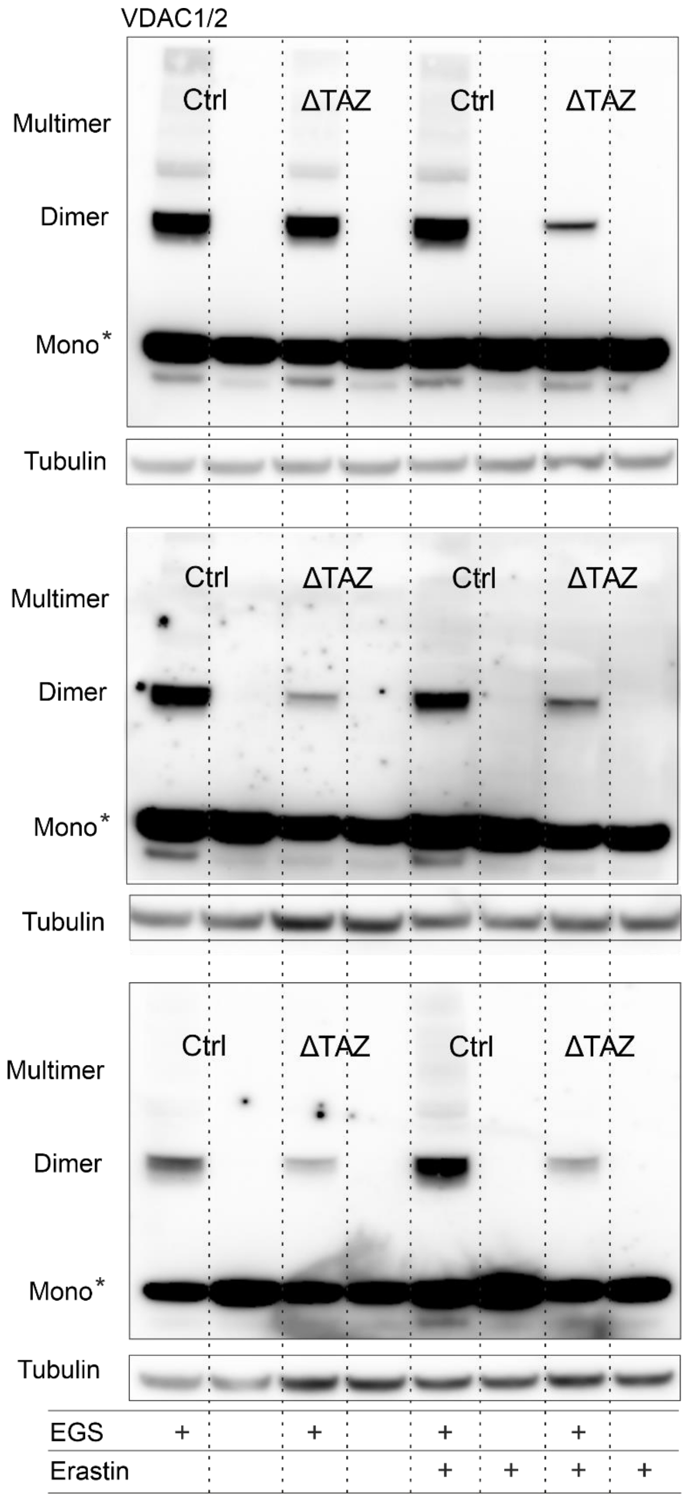


**Supplementary Figure 15: Western blots of VDAC multimer formation upon erastin treatment.** EGS crosslinking was used to assess VDAC multimer levels before and after 8 hours of 15 µM erastin treatment. Ctrl cells exhibited higher levels of VDAC multimers under untreated conditions compared to ΔTAZ cells. Upon erastin treatment, multimer levels were further upregulated in Ctrl cells, whereas ΔTAZ cells failed to show such upregulation. Monomers are oversaturated, indicated by an asterisk (*).
